# Giant ankyrin-B mediates transduction of axon guidance and collateral branch pruning factor Sema 3A

**DOI:** 10.1101/2021.05.03.442401

**Authors:** Blake A. Creighton, Deepa Ajit, Simone Afriyie, Julia Bay, Damaris Lorenzo

**Affiliations:** Department of Cell Biology and Physiology, University of North Carolina at Chapel Hill, Chapel Hill, NC, USA; Carolina Institute for Developmental Disabilities, Chapel Hill, NC, USA; Neuroscience Center, University of North Carolina at Chapel Hill, Chapel Hill, NC, USA

## Abstract

Variants in the high confident autism spectrum disorder gene *ANK2* target both ubiquitously expressed 220-kDa ankyrin-B and neurospecific 440-kDa ankyrin-B (AnkB440) isoforms. Previous work showed that knock-in mice expressing an ASD-linked *Ank2* variant yielding a truncated AnkB440 product exhibit ectopic brain connectivity and behavioral abnormalities. Expression of this variant or loss of AnkB440 caused axonal hyperbranching *in vitro*, which implicated AnkB440 microtubule bundling activity in suppressing collateral branch formation. Leveraging multiple mouse models, cellular assays, and live microscopy, we show that AnkB440 also modulates axon collateral branching stochastically by reducing the number of F-actin-rich branch initiation points. Additionally, we show that AnkB440 enables growth cone (GC) collapse in response to chemorepellent factor semaphorin 3A (Sema 3A) by stabilizing its receptor complex L1 cell adhesion molecule/neuropilin-1. ASD-linked *ANK2* variants failed to rescue Sema 3A-induced GC collapse. We propose that impaired response to repellent cues due to AnkB440 deficits leads to axonal guidance and branch pruning defects and may contribute to the pathogenicity of *ANK2* variants.

## Introduction

Precise wiring of the neural circuitry relies on a tightly regulated spatiotemporal program of axonal extension and pathfinding to form synapses with their targets. The remarkable task of correctly extending the axon through dense three-dimensional spaces such as the developing mammalian brain is driven by the growth cone (GC)–the highly dynamic structure that provides the mechanical force at the tip of the growing axon. To contact targets not in their path and to maximize the number of synapses, most neurons multiply interneuron contact points through collateral axon branches, which are also extended by individual GCs (*Kalil & Dent, 2014; Armijo-Weingart & Gallo, 2017*). In humans, most synapses are assembled during brain development, which number is markedly reduced by adulthood via refinements that include the pruning of axons, collateral branches, dendrites, and dendritic spines (*Riccomagno and Kolodkin, 2015; Low et al., 2008; Petanjek et al., 2011*). The growth and guidance of axonal processes and the sculpting of synaptic connections are determined by multiple intrinsic and extrinsic cellular factors (*Riccomagno and Kolodkin, 2015*). Mistargeting, deficits or excess of axonal processes can lead to hypo-, hyper-, or aberrant neuronal connectivity, which in turn may underlie functional deficits associated with psychiatric and neurodevelopmental disorders (*Holiga et al. 2019; Taquet et al. 2020*).

Ankyrin-B (AnkB) is an integral component of the submembrane cytoskeleton involved in the organization of specialized domains that include ion channels, membranes transporters, cell adhesion molecules (CAMs), and other scaffolding proteins (*Bennett and Lorenzo, 2013; Bennett and Lorenzo, 2016*). AnkB has emerged as a critical determinant of structural neural connectivity through diverse and divergent roles of its two major, alternatively spliced isoforms in the brain; neuronal-specific 440-kDa (“giant”) AnkB (AnkB440) and ubiquitously expressed 220-kDa AnkB (AnkB220) (*Lorenzo, 2020*) (*Figure 1A*). AnkB knockout mice lacking expression of both AnkB220 and AnkB440 exhibit noticeable reduction in the length and number of long axon tracts in multiple brain areas and experience early neonatal mortality (*Scotland et al., 1998; Lorenzo et al., 2014*). Correspondingly, cultured neurons from these mice fail to properly elongate their axons, in part due to reductions in the axonal transport of multiple organelles. AnkB220 enables organelle transport by coupling the retrograde motor complex to vesicles in axons (*Lorenzo et al., 2014*) and other cell types (*Qu et al., 2016; Lorenzo et al., 2015; Lorenzo and Bennett, 2017*). Rescue with AnkB220 is sufficient to restore both normal axonal growth and organelle trafficking *in vitro*. In contrast, the lifespan and overall development of mice lacking only AnkB440 (AnkB440 KO) appear normal. However, AnkB440 KO mice exhibit abnormal communication and sexual behavior responses, repetitive behavior, and increased executive function in the absence of ASD-associated comorbidities (*Yang et al., 2019*). These behavioral deficits are also present in knock-in mice bearing a frameshift mutation (AnkB440^R2589fs^) in the inserted exon unique to AnkB440 that models a human variant found in an individual diagnosed with ASD. AnkB440^R2589fs^ mice have normal AnkB220 expression in the brain and instead of full-length AnkB440 express reduced levels of a truncated 290-kDa AnkB polypeptide lacking a portion of the neuronal-specific domain (NSD) and the entire death and C-terminal regulatory domains (*Yang et al., 2019*). Diffusion tension imaging (DTI) tractography of AnkB440^R2589fs^ brains revealed the presence of ectopic axon projections and stochastic alterations in axonal connectivity, consistent with a potential role of AnkB440 in modulating the formation, guidance, and refinement of axonal connections in the developing brain. In line with these findings, primary cortical neurons from both AnkB440 KO and AnkB440^R2589fs^ mice exhibit increased axonal branching (*Yang et al., 2019*). AnkB440 directly binds and bundles microtubules through a bipartite microtubule-interaction site within the neuronal-specific domain, which is required to suppress ectopic axonal branching in cultured neurons (*Chen et al., 2020*). Furthermore, AnkB440, but not AnkB220, binds L1 cell adhesion molecule (L1CAM) at axonal membranes, where AnkB440 and L1CAM collaborate to coordinate positioning and local dynamics of cortical microtubules and prevent the stabilization of nascent axonal filopodia and maturation into collateral axon branches. In agreement with this model, cultured neurons from knock-in mice bearing the L1CAM variant p.Y1229H, which lacks AnkB440 binding activity, show excess axon collateral branching (*Yang et al., 2019*).

**Figure 1.**
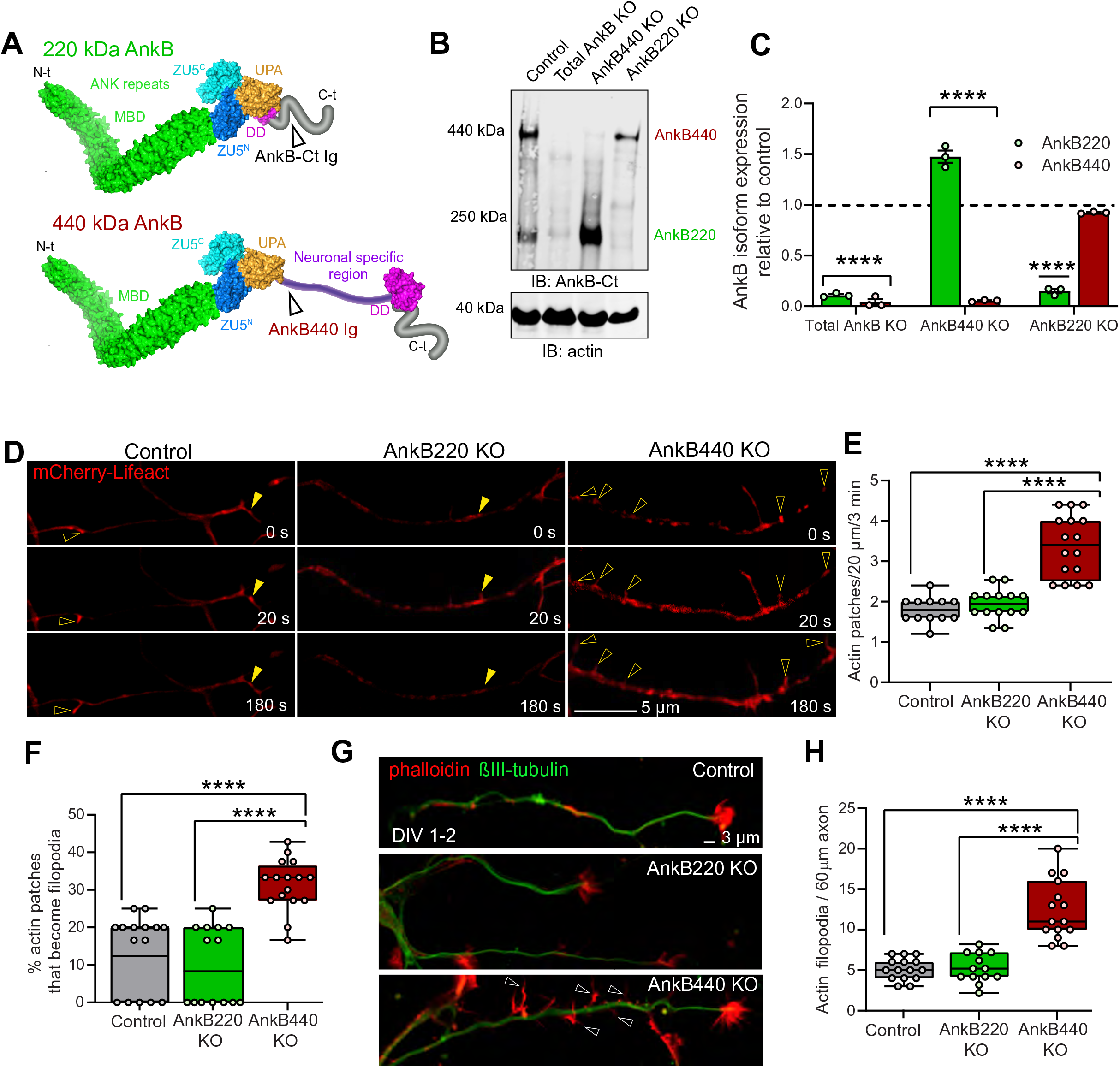
Ankyrin-B 440 suppresses the formation of axonal filopodia *in vitro*. (**A**) Representation of the two major ankyrin-B isoforms in the mammalian brain and their functional domains. The localization of the epitopes recognized by the pan anti-ankyrin B (AnkB-Ct Ig) or the anti 440 kDa ankyrin-B (AnkB440 Ig) antibodies is indicated by arrowheads. (**B**) Western blot analysis of expression of ankyrin-B isoforms in the cortex of PND1 control mice and of mice individually lacking the 220kDa ankyrin-B isoform (AnkB220 KO), the 440 kDa ankyrin-B isoform (AnkB440 KO), and both isoforms (Total AnkB KO). (**C**) Quantification of AnkB220 and AnkB440 levels normalized to actin in cortical lysates from PND1 mice of indicated genotypes relative to their levels in control brains. Data show mean ± SEM for three biological replicates per genotype. Unpaired *t* test. ****p < 0.0001. (**D**) Images selected from timelapse sequences of DIV1-2 neuron expressing mCherry-Lifeact showing the formation of actin patches (asterisks), some of which lead to filopodia that may persist (open arrowheads) or retract (closed arrowheads). Scale bar, 20 μm. (**E**) Quantification of actin patches observed in 20 μm of axons during a 3-minute acquisition period. Data was collected from n=12 control, n=14 AnkB220 KO, n=16 AnkB440 KO axons from three independent experiments. (**F**) Quantification of proportion of actin patches that result in filopodia during the movie acquisition interval derived from analyses of n=15 control, n=14 AnkB220 KO, and n=16 AnkB440 KO axons from three independent experiments. (**G**) Images of DIV1-2 cortical neurons stained for phalloidin and βIII-tubulin. Open arrowheads indicate the presence of axonal filopodia. Scale bar, 3 μm. (**H**) Quantification of filopodia per 60 μm of axons obtained from the analysis of n=15 control, n=13 AnkB220 KO, and n=15 AnkB440 KO axons from three independent experiments. The box and whisker plots in **E**, **F**, and **H** represent all data points collected arranged from minimum to maximum. One-way ANOVA with Dunnett’s post hoc analysis test for multiple comparisons. ****p < 0.0001.

Here we show that AnkB440 also modulates axon collateral branching stochastically by suppressing the formation of F-actin-rich branch initiation points. We demonstrate that axons require AnkB440, but not AnkB220, to transduce the chemorepellent signals induced by the extrinsic factor semaphorin-3A (Sema 3A). GCs in the main axon and in collateral branches of cultured cortical neurons from AnkB440 KO mice fail to collapse in response to Sema 3A. This suggests that in addition to suppressing axon collateral branch initiation, AnkB440 is involved in collateral branch pruning. However, unlike the suppression of branch point initiation (*Chen et al., 2020*), GC collapse and branch retraction did not involve the interaction of AnkB440 with microtubules. On the other hand, binding activity between AnkB440 and L1CAM is necessary for normal transduction of Sema 3A cues and for downstream actin cytoskeleton remodeling, but it does not involve AnkB partner βII-spectrin. Lastly, we found that selected *ANK2* variants found in individuals with ASD phenocopied the loss of response to Sema 3A, indicating a potential mechanism of pathogenicity.

## Results

### Loss of AnkB440, but not AnkB220, results in higher density of actin patches, filopodia, and collateral axon branches

Most collateral branches emerge from transient actin-rich nucleation sites along the axon, which give rise to branch precursor filopodia (*Gallo, 2011*). Commitment to branch formation requires microtubule invasion of the nascent filopodia (*Yu et al., 1994; Kalil & Dent, 2014*). AnkB440 recruits and stabilizes microtubules at the plasma membrane and its loss has been proposed to cause axonal hyperbranching *in vitro* by promoting microtubule unbundling and invasion of nascent branches (*Yang et al., 2019; Chen et al., 2020*). AnkB440 loss results in 50% upregulation of AnkB220 in brains of AnkB440 KO mice (*Yang et al., 2019*) (*Figure 1B,C;* see *Figure 1-source data 1-3*). We previously reported that AnkB220 promotes axonal growth and axonal transport of organelles, including mitochondria (*Lorenzo et al., 2014*), which has been implicated in axonal hyperbranching (*Courchet et al., 2013*). Likewise, AnkB220 associates with microtubules in biochemical assays (*Davis and Bennett, 1984*). Changes in AnkB220 expression could contribute to or compensate for deficits in AnkB440, and its effects on axonal branching. Thus, we sought to determine whether AnkB220 and AnkB440 have independent roles in axonal branch formation using cortical neurons harvested from AnkB440 KO mice (*Yang et al., 2019*; *Chen et al., 2020*) and from a novel conditional AnkB220 KO mouse model (herein referred as AnkB220 KO) (*Figure 1B,* see *Figure 1-source data 1-3*). The conditional *Ank2* allele used to selectively eliminate AnkB220 expression was generated by fusing exons 36 and 37 of *Ank2* to prevent splicing of exon 37 unique to the AnkB440 transcript (*Figure 1—figure supplement 1A*). Western blot analysis from cortical lysates of mice in which recombination of the *Ank2* floxed allele (*Figure 1—figure supplement 1B, see Figure 1-supp Fig 1 source data 1,2*) was induced by Nestin-Cre (*AnkB220^flox/flox^*/*Nestin-Cre;* AnkB220 KO) demonstrates efficient knockdown of AnkB220 and preservation of AnkB440 expression (*Figure 1A,B*).

The focal accumulation of dynamic pools of F-actin, termed actin patches, along the axon shaft have been shown to promote filopodia formation, the first step during collateral axon branch development (*Gallo, 2011; Spillane et al., 2011; Armijo-Weingart and Gallo; 2017*). Furthermore, the presence of larger and more stable actin patches within the primary axon directly correlates with the formation of axon protrusions *in vivo* (*Hand et al., 2015)*. We found that loss of AnkB440 in day *in vitro* 1 (DIV1) cortical neurons resulted in increased number of actin patches relative to control and AnkB220 KO neurons (*Figure 1D,E, arrowheads*). These mCherry-Lifeact-labeled F-actin patches marked sites for emerging filopodia that either continued to extend (*Figure 1D, open arrowheads*) or retracted and vanished (*Figure 1D, closed arrowheads*). We found that a higher proportion of actin patches led to the emergence of filopodia protrusions in AnkB440 KO neurons (*Figure 1F*), which correlated with a higher density of both axonal filopodia (*Figure 1G,H*) and of collateral axon branches (*Figure 1—figure supplement 2A,D*) relative to control and AnkB220 KO neurons. Loss of AnkB220, but not of AnkB440, resulted in shorter axons (*Figure 1—figure supplement 2A-C*) and impaired organelle transport (*Figure 1—figure supplement 3A-D*) (*Figure 1—video 1*), which confirms our previous findings based on *in vitro* rescue experiments that AnkB220 is critical for axonal growth and organelle dynamics (*Lorenzo et al., 2014*). This indicates that while AnkB440 suppresses collateral branch formation, AnkB220 is not a cell-autonomous determinant of axonal branching, and that elevations in AnkB220 do not influence organelle dynamics in AnkB440 deficient axons. Together, our data suggest that in addition to the previously proposed role of loss of AnkB440 in stabilizing axon branching by promoting microtubule invasion of nascent axonal filopodia (*Yang et al., 2019*), AnkB440 deficits also contribute to axonal hyperbranching stochastically by inducing the formation of higher number of axonal filopodia.

### Loss of AnkB440 results in corpus callosum thickening and increased axonal collateral branching *in vivo*

DTI tractography analysis of AnkB440^R2589fs^ brains, which lack full-length AnkB440 but express a truncated 290 kDa AnkB440 polypeptide, show normal organization of major axon tracts, including the corpus callosum (CC) (*Yang et al., 2019*). However, expression of truncated AnkB440 failed to suppress collateral axon branching *in vitro* and caused stochastic increases in cortical axon connectivity *in vivo* (*Yang et al., 2019*). Unlike AnkB440 that preferentially localizes to axons, the truncated fragment is equally distributed between dendrites and axons (*Yang et al., 2019*). The changes in distribution of this AnkB440 fragment, which retains some of the domains and functions of AnkB440, could lead to gain-of-function or dominant negative effects that may differentially influence structural or functional axonal connectivity relative to total AnkB440 loss. We found that postnatal day 25 (PND25) AnkB440 KO mice exhibit CC hyperplasia (*Figure 2A,B*). Increased CC thickness in AnkB440 KO brains is not due to changes in the abundance and distribution of callosal projection neurons in neocortical layers, assessed by expression of the transcription factors Satb2, Ctip2 and Tbr1 (*Fame at al., 2011*) (*Figure 2C,D)*. To determine whether volumetric gains in CC thickness in AnkB440 KO brains could result from axon hyperbranching, we visualized projection neurons from upper layers of the primary somatosensory cortex of PND25 brains using Golgi staining. In agreement with the *in vitro* results (*Figure 1—figure supplement 2A,D*), AnkB440 KO axons traced from brain slices had on average four times more branches than both AnkB220 KO and control axons (*Figure 2D,E*). These increases in axon branching observed in AnkB440 KO brains may alter local and long-range axonal connectivity as observed in AnkB440^R2589fs^ mice (*Yang et al., 2019*).

**Figure 2.**
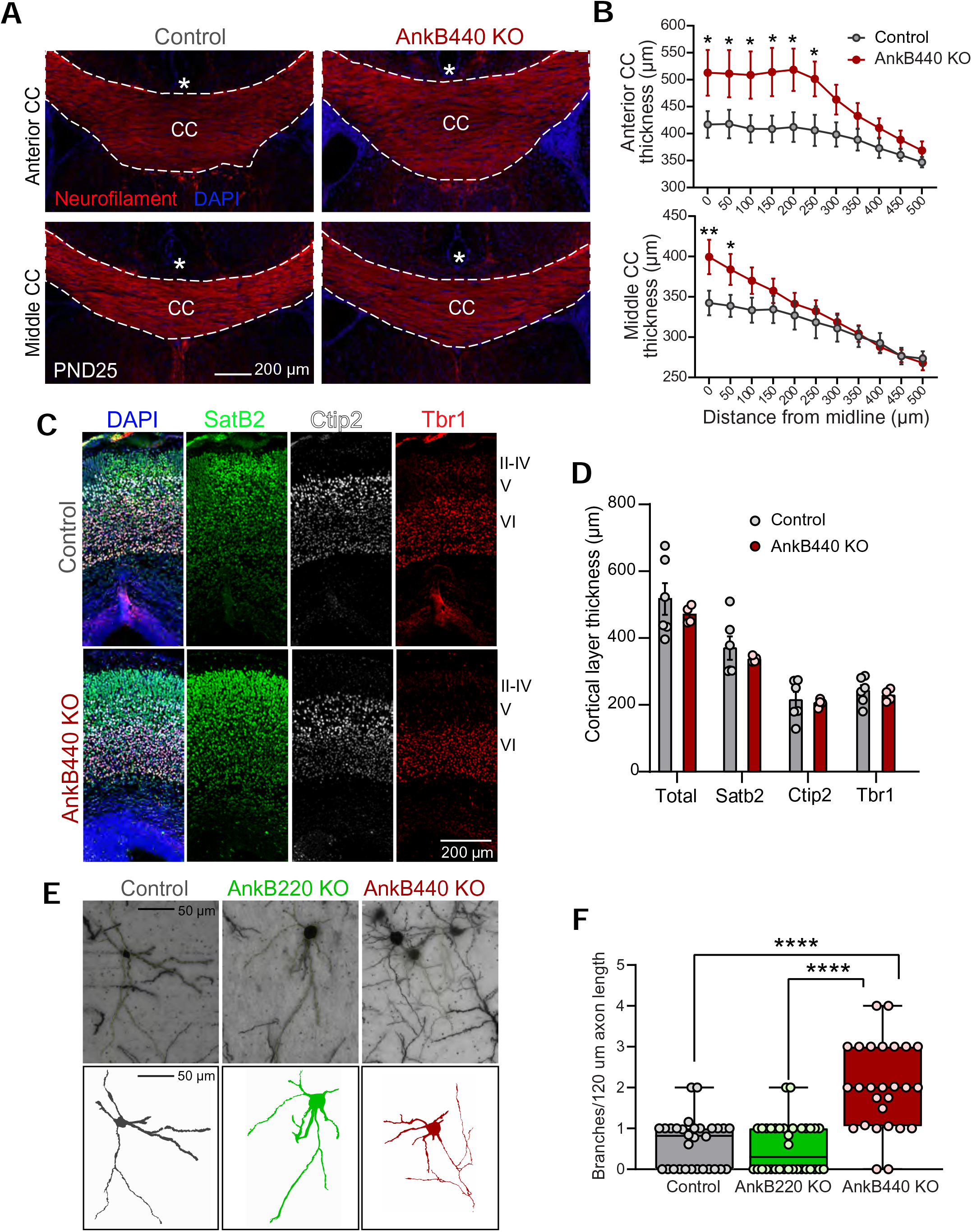
Loss of AnkB440 results in callosal hyperplasia and axon hyperbranching *in vivo*. (**A**) Images of coronal sections of PND25 mouse brains taken through the anterior and middle portions of the corpus callosum (CC) and stained for neurofilament to label axons and DAPI to stain nuclei. An asterisk indicates the position of the brain midline. Scale bar, 200 μm. (**B**) Quantification of thickness of anterior (top) and middle (bottom) regions of the CC at different points from the brain midline from control (n=4) and AnkB440 KO (n=5) brains. (**C**) Images of PND0 brains stained for Satb2, Ctip2, and Tbr1 to label neocortical layers and DAPI to stain nuclei. Scale bar, 200 μm. (**D**) Quantification of total, Sabt2-, Ctip2-, and Tbr1-positive cortical layer thickness cortical thickness from PND0 control (n=5-6) and AnkB440 KO (n=4) brains. Data in **B** and **D** represent mean ± SEM. Unpaired *t* test. **p < 0.01, *p < 0.05. (**E**) Images of Golgi-stained (top) projection neurons in layers II-III of the primary somatosensory cortex of PND25 mice and corresponding traces (bottom). Scale bar, 50 μm. (**F**) Quantification of axonal branches per 120 μm of axon (n=30 control, n=34 AnkB220 KO, and n=27 AnkB440 KO) Golgi-stained neurons. The box and whisker plots represent all data points collected arranged from minimum to maximum. One-way ANOVA with Dunnett’s post hoc analysis test for multiple comparisons. ****p < 0.0001.

### AnkB440 is enriched at axonal growth cones and required for normal responses to Semaphorin 3A

Previous studies suggest that AnkB440 preferentially localizes to axons (*Chan et al., 1993; Yang et al., 2019; Chen et al., 2020*). Staining of DIV3 cortical neurons with an antibody specific for AnkB440 (*Yang et al., 2019*) (*Figure 3A,* see *Figure 3-source data 1-3*) confirmed the widespread AnkB440 distribution in axons (*Figure 3B*), with strong AnkB440 signals also detected at the tips of axonal process, including nascent filopodia, and at the growth cones (GCs) of the main axon and of collateral axon branches (*Figure 3B,C*). As expected, AnkB440 signal was absent from axons of AnkB440 KO neurons stained with the AnkB440 antibody (*Figure 3Di*). Neither the overall axonal distribution nor the localization of AnkB440 at GCs changed in neurons selectively lacking AnkB220 stained with AnkB440-specific (*Figure 3Dii*) or pan-AnkB (*Figure 3Diii*) antibodies. Interestingly, AnkB440 localized to both microtubule- and F-actin-enriched axonal domains (*Figure 3C,D*), respectively stained with βIII-tubulin and phalloidin, suggesting that it might promote the organization and/or dynamics of both cytoskeletal networks.

**Figure 3.**
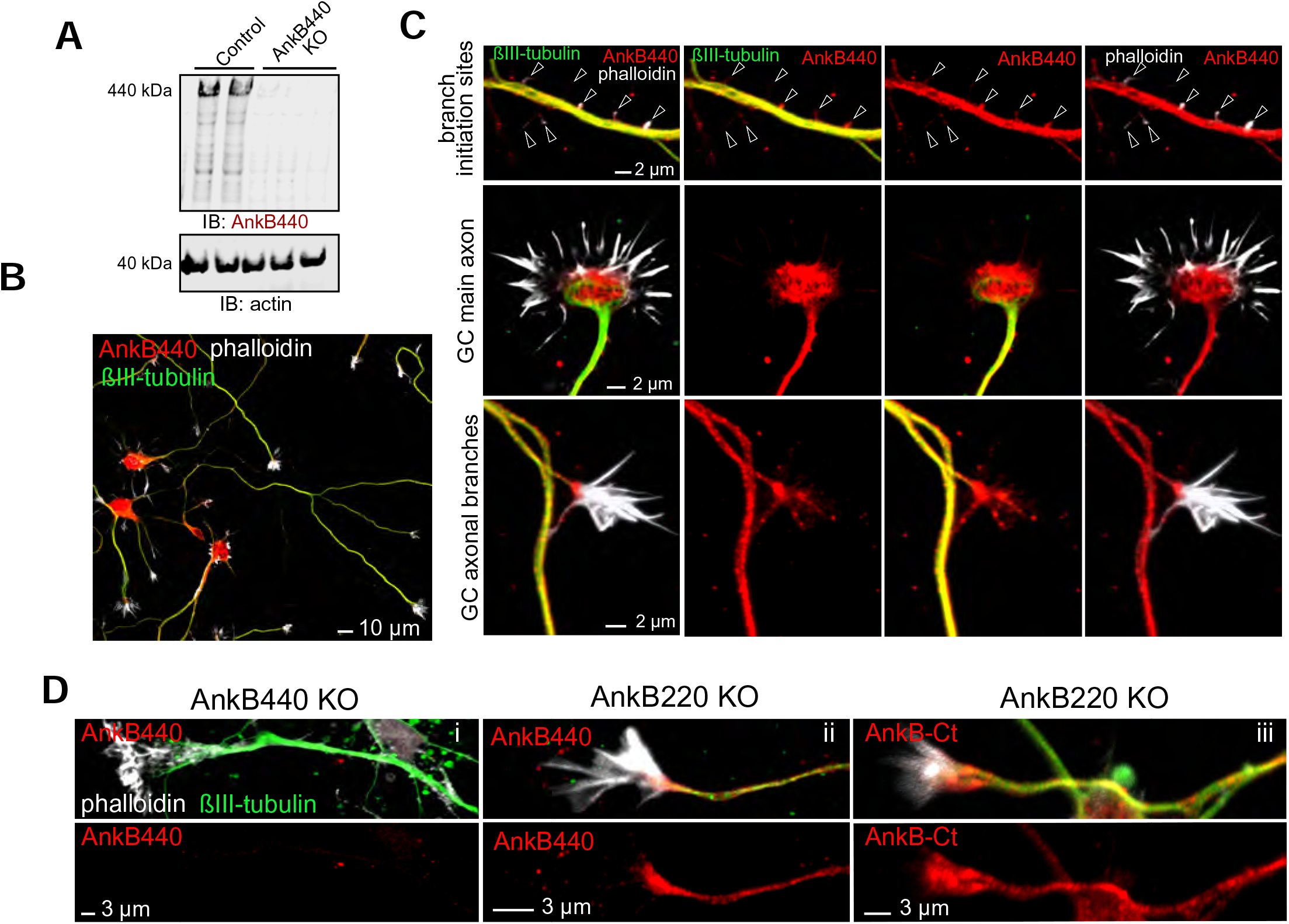
AnkB440 is enriched at axonal growth cones. (**A**) Western blot analysis of expression of AnkB440 in total cortical lysates of PND1 control and AnkB440 KO mice show specificity of the antibody against the AnkB440 isoform. (**B**) Images of DIV3 control cortical neurons stained with the AnkB440-specific antibody, phalloidin (to label F-actin), and βIII-tubulin indicate the broad distribution of AnkB440 in axons. Scale bar, 10 μm. (**C**) Higher magnification images show that AnkB440 is enriched at sites of nascent axonal filopodia (arrowheads, already present before microtubule invasion), and at the growth cone (GC) of the main axon and of collateral axon branches. Scale bar, 2 μm. AnkB440 staining colocalizes with both F-actin and microtubule enriched axonal structures. (**D**) (i) DIV3 AnkB440 KO cortical neurons show specificity of the antibody against the AnkB440 isoform. Staining of AnkB220 KO neurons with AnkB440-specific (ii) and AnkB-Ct (iii) antibodies show the localization of AnkB440 in the axon and its accumulation at the GC. Scale bar, 3 μm.

AnkB440 interacts with L1CAM (*Yang et al., 2019*), which is required for transducing the repulsive growth and guidance effects of the soluble guidance cue class III semaphorin A (Sema 3A) (*Castellani et al., 2000; Castellani et al., 2002*; *Castellani et al.,* 2004). Sema 3A promotes GC collapse (*Luo et al., 1993; Kolodkin et al., 1993*) and inhibits axon branching *in vitro* (*Dent et al., 2004*), which likely underlies its roles in repelling cortical axons and in pruning axonal branches *in vivo* and *in vitro* (*Polleux et al., 1998; Bagri et al., 2003*). Thus, we next evaluated the response of AnkB deficient cortical neurons to Sema 3A. We observed that roughly 60% of GCs of both control and AnkB220 KO cortical neurons collapsed after 30 minutes of Sema 3A treatment, up from a baseline of 20% GC collapse in untreated cells, without any appreciable difference between the two groups (*Figure 4A,B*). In contrast, GCs of total AnkB KO and AnkB440 KO neurons significantly failed to collapse when exposed to Sema 3A (*Figure 4A,C,D*). The failure of GCs to collapse upon Sema 3A treatment in AnkB440 KO neurons was restored by transfection with AnkB440 but not with AnkB220 plasmids (*Figure 4E-G*). To rule out that the inability of Sema 3A to collapse GCs in AnkB440 KO neurons was due to a generalized loss of transduction of chemorepulsive cues, we treated control and AnkB440 neurons with Ephrin A5, which also induces GC collapse in cortical neurons (*Meima et al., 1999*). Ephrin A5 promoted GC collapse in AnkB440 KO cultured cortical neurons at levels indistinguishable from control neurons (*Figure 4—figure supplement 4*). These results indicate a selective requirement for AnkB440 for cortical neuron responses to the GC collapse-inducing effect of Sema 3A.

**Figure 4.**
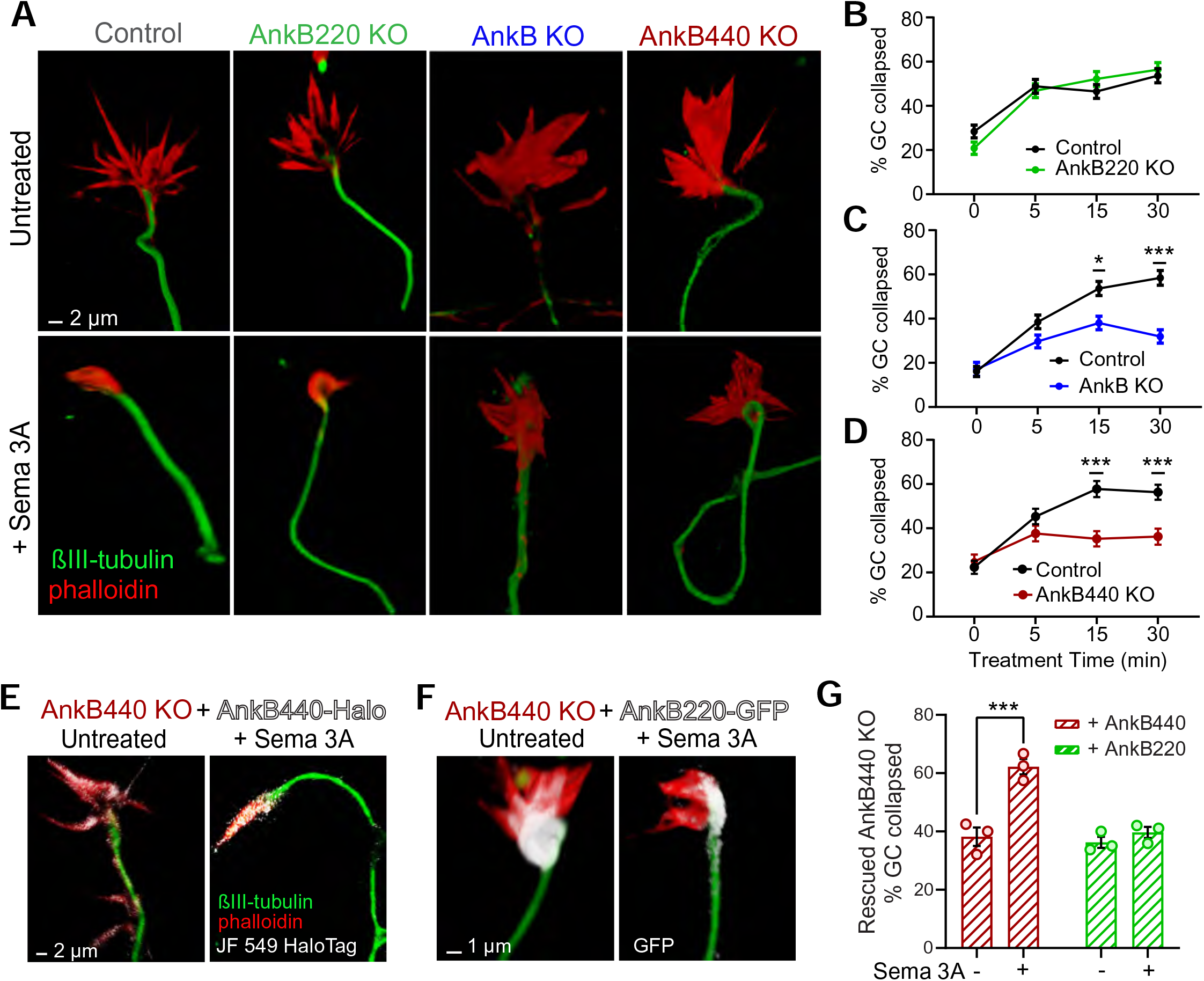
AnkB440 is required for semaphorin 3A-induced GC collapse. (**A**) Images of the distal portion of axons of DIV3 cortical neurons untreated and treated with semaphorin 3A (Sema 3A) and stained with phalloidin and βIII-tubulin. Scale bar, 2 μm. (**B-D**) Percent of GC collapsed before and after Sema 3A treatment of control, AnkB220 (**B**), total AnkB KO (**C**) and AnkB440 KO (**D**) cortical neurons. Data in B-D represent mean ± SEM collected from an average of n=120-200 GCs/treatment condition/genotype from three independent experiments. Unpaired *t* test. ***p < 0.001, *p < 0.05. (**E**) Images of DIV3 AnkB440 KO cortical neurons expressing Halo-tagged AnkB440 untreated and treated with Sema 3A and stained with Janelia Fluor 549 HaloTag ligand to visualize AnkB440. Scale bar, 2 μm. (**F**) Images of DIV3 AnkB440 KO cortical neurons expressing GFP-tagged AnkB220 untreated and treated with Sema 3A and stained for GFP to visualize AnkB220. Scale bar, 1 μm. (**G**) Percent of GC collapsed before and after Sema 3A treatment of AnkB440 KO neurons rescued with AnkB440 or AnkB220 cDNAs. Data represent mean ± SEM collected from an average of n=150 GCs/treatment condition/transfection from three independent experiments. Unpaired *t* test. ***p < 0.001.

### The AnkB440-L1CAM complex promotes GC collapse in response to Sema 3A

To confirm that AnkB440 and L1CAM associate at GCs of cultured neurons, we performed a proximity ligation assay (PLA) using antibodies specific for L1CAM and AnkB440, which produces a signal when both proteins are within 40 nm of each other (*Fredriksson et al., 2002; Lorenzo and Bennett, 2017*). PLA signal was detected at the GC and throughout the axons in control neurons, but was virtually lost in AnkB440 KO neurons and in L1CAM neurons harboring the p.Y1229H variant (*Figure 5A*), which lacks AnkB440 binding activity, consistent with previous findings *in vivo* and *in vitro* (*Yang et al., 2019*). We further confirmed the specific interaction of L1CAM with AnkB440 using a cellular assay where co-expression of each GFP-tagged AnkB isoform with L1CAM in HEK293T cells resulted in the effective recruitment of GFP-AnkB440, but not of GFP-AnkB220, from the cytoplasm to the plasma membrane (*Figure 5—figure supplement 5*). Notably, treatment of p.Y1229H L1CAM neurons with Sema 3A resulted in negligible changes in the percent of collapsed GCs in both the axon and collateral axonal branches (*Figure 5B-E*. Loss of collateral axon branch GC collapse was also detected by live imaging in AnkB440 KO neurons (*Figure 5D,E*) (*Figure 5—video 2*), which could contribute to the aberrant development and connectivity of major axonal tracts, and to the hyperplasia of the CC detected in AnkB440 KO mice (*Figure 2A,B*). Taken together, these results support a critical role for AnkB440 and the AnkB440-L1CAM complex in Sema 3A-induced GC collapse.

**Figure 5.**
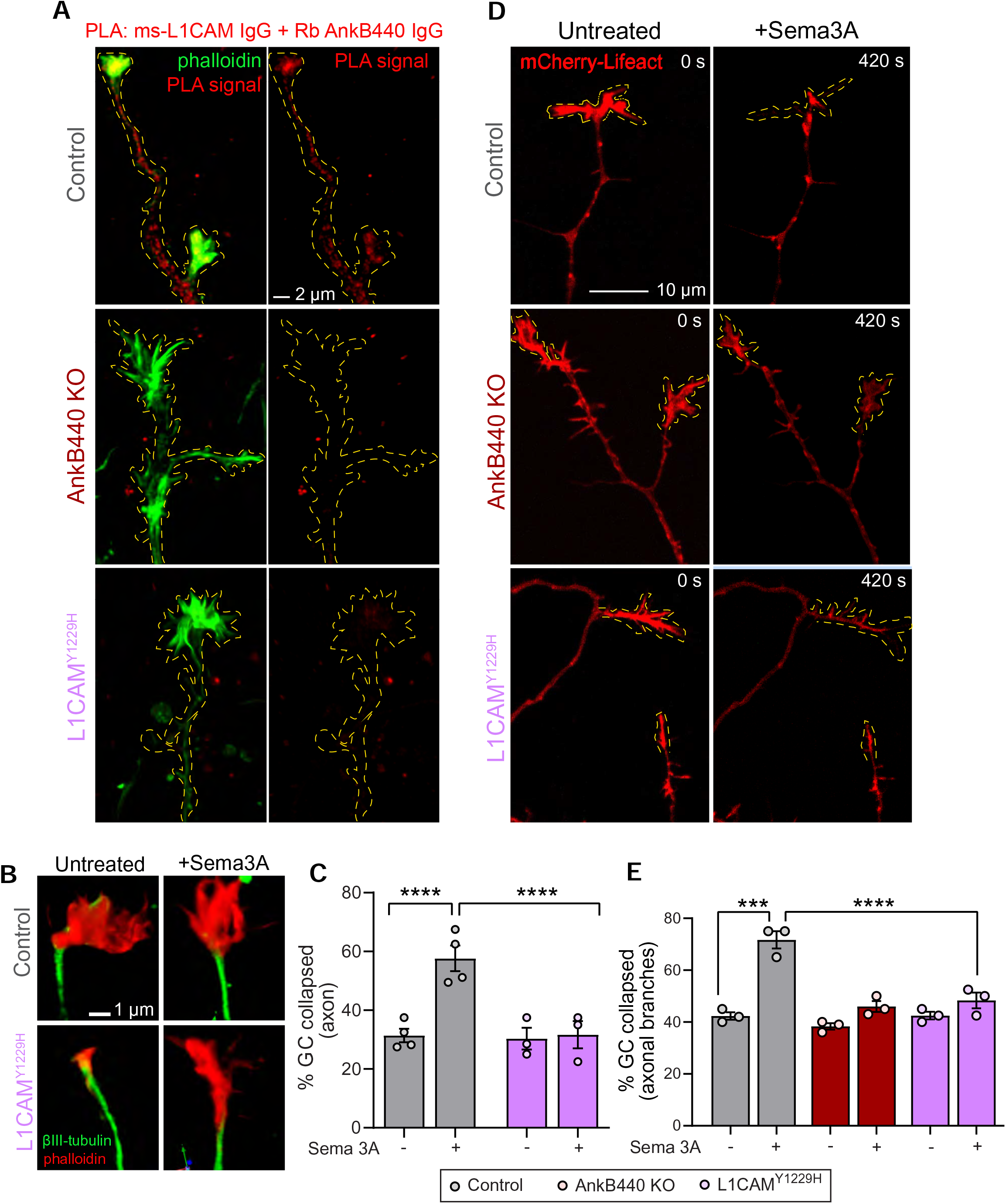
The AnkB440-L1CAM complex promotes GC collapse induced by Sema 3A. (**A**) Images show PLA signal between AnkB440 and L1CAM in axons and GCs of DIV3 cortical neurons from the indicated genotypes. Phalloidin staining was used to identify GCs. Scale bar, 2 μm. (**B**) Images of the distal portion of the main axon untreated and treated with Sema 3A and stained with phalloidin and βIII-tubulin. Scale bar, 1 μm. (**C**) Percent of axon GCs that collapse before and after Sema 3A treatment. (**D**) Images from timelapse sequences of DIV3 cortical neuron expressing mCherry-Lifeact before and after Sema 3A treatment. Scale bar, 10 μm. (**E**) Percentage of collateral axon branches GCs that collapse before and after Sema 3A treatment. Data in **C** and **E** represent mean ± SEM collected from an average of n=80 GCs/treatment condition/genotype from three independent experiments. One-way ANOVA with Dunnett’s post hoc analysis test for multiple comparisons. ****p < 0.0001, ***p < 0.001.

### AnkB440 stabilizes L1CAM and the Sema 3A receptor Nrp1 at the cell surface of GCs

We next sought to identify the mechanisms by which AnkB440 and its interaction with L1CAM promotes GC collapse in response to Sema 3A. First, we established that AnkB440 loss does not alter L1CAM expression in total lysates from PND1 AnkB440 KO cortex or from AnkB440 KO cortical neuron cultures (*Figure 6A-D*, see *Figure 6-source data 1-10)*. Using a biotinylation assay that labels and captures surface proteins in cortical neuron cultures, we determined that AnkB440 loss markedly reduced surface levels of L1CAM relative to its membrane abundance in control neurons, while surface levels of the AMPA receptor subunit GluR1 remained unchanged (*Figure 6C,D, Figure 6-source data 1-10*). These findings indicate that AnkB440 is required to maintain normal levels of L1CAM at the cell surface of neurons. L1CAM is not a Sema 3A receptor (*Castellani et al., 2000*). Instead, L1CAM directly binds the Sema 3A receptor neuropilin-1 (Nrp1) (*Chen et al., 1998*), which due to its short cytoplasmic domain requires the formation of complexes with transmembrane co-receptors to be stably anchored at the cell surface and propagate repulsive Sema 3A signals (*Castellani et al., 2000; Castellani et al., 2002*). In addition to the L1CAM-Nrp1 complex, Plexin A1 also takes part in the transduction of Sema 3A signals (*Bechara et al., 2008*). Thus, we next evaluated whether these two receptors participate in AnkB440-L1CAM modulation of Sema 3A effects. Loss of AnkB440 KO led to a slight increase of Nrp1 levels in total cortical lysates (*Figure 6A,B*) and to a marked reduction of surface-bound Nrp1 in AnkB440 KO cortical neurons, assessed via surface biotinylation, despite its increased total expression (*Figure 6C,D*). In contrast, Plexin A1 levels remained unchanged both in AnkB440 KO brains (*Figure 6A,B*) and at the surface of AnkB440 KO neurons (*Figure 6C,D*). Noticeably, Sema 3A levels were increased in the cortex of AnkB440 KO PND1 mice (*Figure 6A,B*), which together with the smaller increase in Nrp1 (*Figure 6A-D*) points towards protein upregulation as a likely mechanism to compensate for the observed loss of Sema 3A-mediated GC collapse responses.

**Figure 6.**
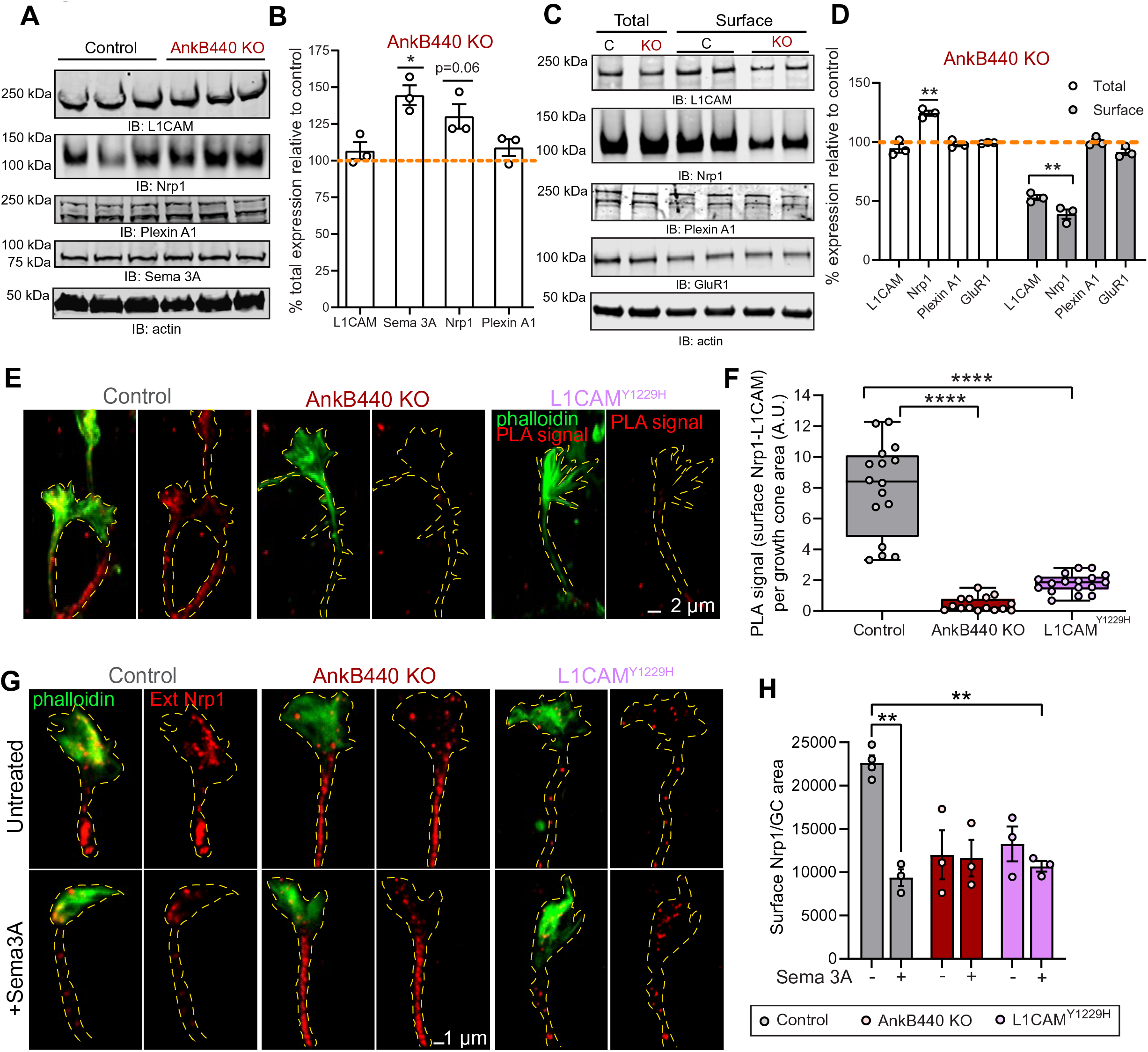
AnkB440 stabilizes the Sema 3A receptor complex L1CAM-Nrp1 at the cell surface of GCs. (**A**) Western blot analysis of the expression of Sema 3A, L1CAM and the Sema 3A receptors Nrp1 and Plexin A1 in the cortex of PND1 control and AnkB440 KO mice. (**B**) Quantification of protein levels normalized to actin in cortical lysates from PND1 mice of indicated genotypes relative to their levels in control brains. Data show mean ± SEM for three biological replicates per genotype. Unpaired *t* test. *p < 0.05. (**C**) Western blot analysis of total and surface levels of indicated proteins in DIV3 control and AnkB440 KO cortical neurons. (**D**) Quantification of total and surface protein levels normalized to actin relative to their levels in control cortical neurons. Data show mean ± SEM for three biological replicates per genotype. Unpaired *t* test. **p < 0.01. (**E**) Images show PLA signal between L1CAM in Nrp1 at the cell surface of the axon and GCs of DIV3 cortical neurons from the indicated genotypes. This assay used an Nrp1 antibody that selectively recognizes and extracellular epitope. Phalloidin staining was used to identify GCs. Scale bar, 2 μm. (**F**) Quantification of PLA signal at GC surface relative to GC area collected from an average n=15 GCs/genotype. The box and whisker plots represent all data points collected arranged from minimum to maximum. One-way ANOVA with Dunnett’s post hoc analysis test for multiple comparisons. ****p < 0.0001. (**G**) Images show Nrp1 localization at the surface of GC of DIV3 neurons untreated and treated with Sema 3A, which induces the internalization of surface Nrp1. Scale bar, 1 μm. (**H**) Quantification of surface Nrp1 levels at GCs relative to GC area at the basal state and upon Sema 3A-induced Nrp1 internalization. Data represent mean ± SEM collected from an average of n=90 GCs/treatment condition/genotype from three independent experiments. One-way ANOVA with Dunnett’s post hoc analysis test for multiple comparisons. **p < 0.01.

To confirm whether surface levels of L1CAM-Nrp1 complexes were altered in GCs of AnkB440 KO cortical neurons, we selectively labeled surface Nrp1 in non-permeabilized DIV3 neurons with an Nrp1 antibody that recognizes an extracellular epitope. Then, we labeled all L1CAM molecules upon cell permeabilization and visualized surface Nrp1-L1CAM complexes using the PLA assay. PLA signal between surface Nrp1 and L1CAM, indicative of surface Nrp1-L1CAM complexes, was abundant at the GC and in the axon of control neurons, but significantly reduced in AnkB440 KO neurons (*Figure 6E,F*). Consistent with the proposed role of AnkB440 in promoting L1CAM, and consequently, Nrp1 localization at the cell surface, p.Y1229H L1CAM neurons in which the AnkB440-L1CAM association is disrupted, also showed a dramatic reduction in surface L1CAM-Nrp1 PLA signal (*Figure 6E,F*). Both AnkB440 KO and p.Y1229H L1CAM neurons exhibited reduced internalization of surface Nrp1 in response to Sema 3A, consistent with previous reports showing that Nrp1 internalization is required during Sema 3A-induced GC collapse (*Castellani et al., 2004*). Together, these results indicate that AnkB440 promotes the membrane surface localization of the L1CAM-Nrp1 holoreceptor complex, which is required to respond to Sema 3A repulsive cues.

### AnkB440 does not require βII-spectrin or binding to microtubules to transduce Sema 3A signals during GC collapse

AnkB directly binds βII-spectrin (*Davis and Bennett, 1984; Davis et al., 2009*), which is widely distributed along axons and required for axonal elongation and organelle transport (*Lorenzo et al., 2019*). βII-spectrin binds F-actin and tetramers of βII-spectrin/αII-spectrin organize a periodic submembrane network of F-actin and associated proteins throughout all axonal domains (*Xu et al., 2013*). Thus, we next investigated whether βII-spectrin is required to propagate Sema 3A signals to the F-actin network during GC collapse. GCs of cortical neurons harvested from mice lacking βII-spectrin in the brain (βII-spectrin KO) (*Galiano et al., 2012; Lorenzo et al., 2019*) collapsed normally in response to Sema 3A treatment (*Figure 6—figure supplement 6A,B*). This result indicates that AnkB440 promotes Sema 3A-induced GC collapse independently of βII-spectrin.

AnkB440 also binds and bundles microtubules through a bipartite microtubule-interaction site located in its NSD (*Chen et al., 2020*). This microtubule-binding activity is required to suppress ectopic axon branching in cultured neurons *Chen et al., 2020*). The site of microtubule interaction in AnkB440 comprises a module of 15 tandem imperfect 12-aa repeats that includes highly conserved residues in the third (Pro, P), fifth (Ser, S), and ninth positions (Lys, K) (*Chen et al., 2020*). Point mutations in each of these residues in full length AnkB440 (PSK mutant) is sufficient to impair microtubule-binding activity and cause axon hyperbranching *in vitro* (*Chen et al., 2020*). To test whether AnkB440-microtubule binding is required for Sema 3A-induced GC collapse, we transfected Halo-tagged cDNA of AnkB440-PSK into AnkB440 KO cortical neurons and treated them with Sema 3A. Like in control neurons, about 60% of GCs of neurons expressing AnkB440-PSK collapsed in response to Sema 3A, indicating that the AnkB440-microtubule interaction is not required for normal transduction of Sema 3A signals (*Figure 6—figure supplement 6C,D*).

### AnkB440 and its interaction with L1CAM are required to modulate cofilin activity in response to Sema 3A

GC motility and collapse involves fast turnover and reorganization of actin filaments (*Omotade et al., 2017*). F-actin dynamics is regulated by the actin depolymerizing and severing factor (ADF)/cofilin (*Carlier et al., 1997; Maciver, 1998*). Cofilin phosphorylation by the Ser/Thr kinase LIM-kinase (LIMK) at the Ser3 site (*Arber et al., 1998; Yang et al., 1998*) inactivates cofilin and prevents its F-actin severing activity (*Agnew et al., 1995; Moriyama et al., 1995).* Similarly, LIMK phosphorylation of cofilin and the rapid subsequent cofilin activation have been shown to be critical steps in the disassembly of actin filaments during Sema 3A-induced GC collapse (*Aizawa et al., 2001*). Thus, we evaluated whether loss of AnkB440 led to changes in expression of LIMK and cofilin, or in cofilin phosphorylation. We found that total levels of LIMK and cofilin were not altered in cortical lysates from PND1 AnkB440 brains (*Figure 7A,B, see Figure 7-source data 1-5*). However, AnkB440 loss decreased the ratio of inactive phospho-cofilin (pcofilin^S3^) to total cofilin by 40% relative to control (*Figure 7A,B, see Figure 7-source data 1-5*). This increase in active cofilin in AnkB440 mice during early brain development might provide a more dynamic actin pool and could underlie the surges in axonal actin patches and emerging filopodia observed in AnkB440 KO cortical neurons.

**Figure 7.**
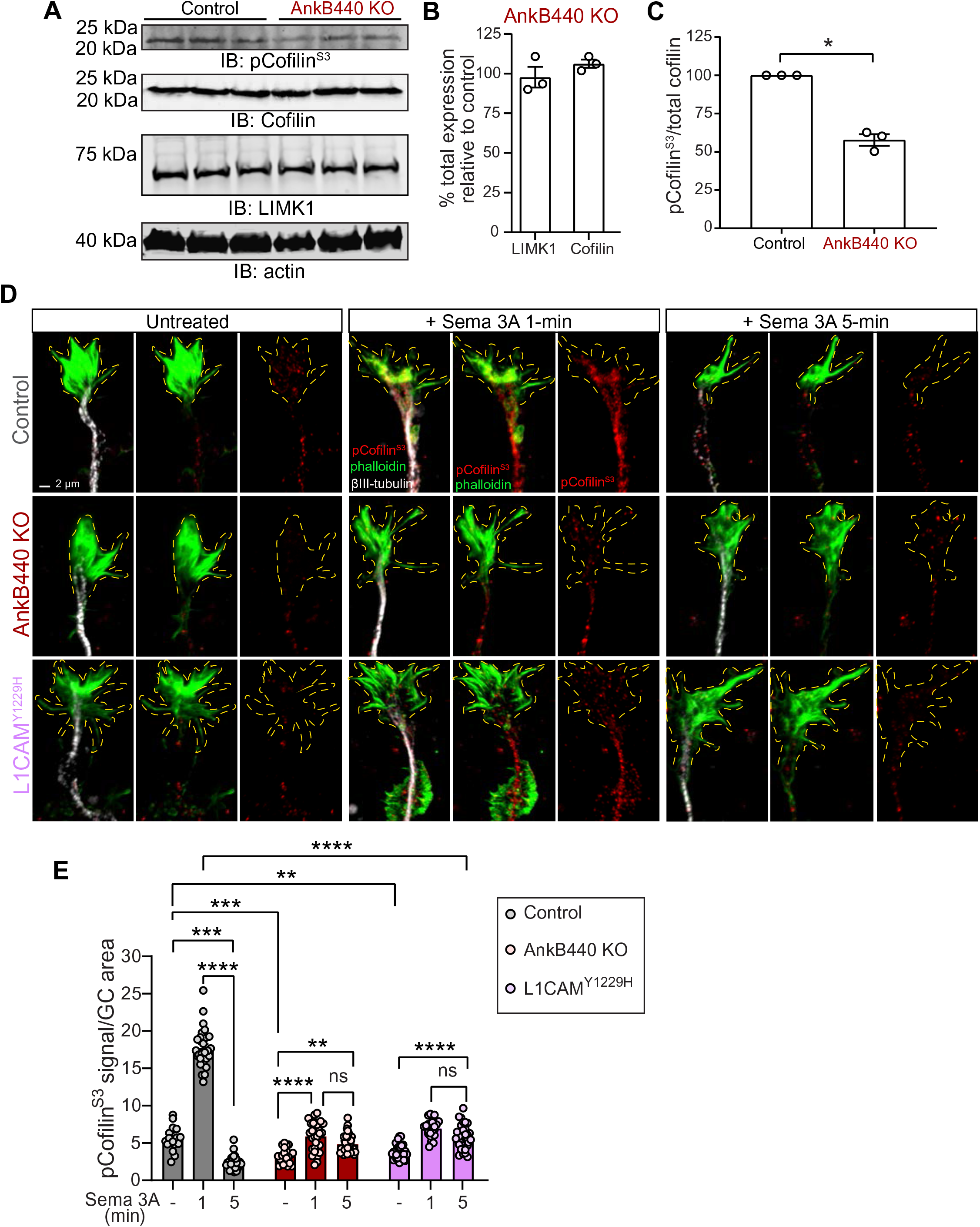
AnkB440 and its interaction with L1CAM are required for F-actin disassembly during GC collapse upon Sema 3A signaling. (**A**) (Western blot analysis of the expression of Cofilin, phospho-cofilin (Ser3) and LIMK1 in the cortex of PND1 control and AnkB440 KO mice. Actin is a loading control. (**B**) Quantification of total levels of cofilin and LIMK1 normalized to actin in cortical lysates from PND1 AnkB440 KO mice relative to their levels in control brains. (**C**) Quantification of levels of phosho-cofilin relative to total cofilin in cortical lysates from PND1 control and AnkB440 KO mice. Data in **B** and **C** represent mean ± SEM for three biological replicates per genotype. Unpaired *t* test. *p < 0.05. (**D**) Images of the distal portion of the main axon of DIV3 cortical neurons untreated and treated with Sema 3A for 1 and 5 minutes and stained with phalloidin, βIII-tubulin, and pCofilin^Ser3^. Dotted lines indicate GCs. Scale bar, 2 μm. (**E**) Quantification of pCofilin^S3^ signal at GCs relative to GC area at the basal state and upon Sema 3A treatment for 1 and 5 minutes. Data represent mean ± SEM collected from an average of n=25-40 GCs/treatment condition/genotype from three independent experiments. One-way ANOVA with Dunnett’s post hoc analysis test for multiple comparisons. ****p < 0.0001, ***p < 0.001, **p < 0.01.

To determine whether AnkB440 participates in Sema-3A-induced regulation of actin dynamics through the action of cofilin, we examined the effect of Sema-3A on the localized activation/inactivation of cofilin at GCs through confocal microscopy. As previously observed in cultured dorsal root ganglion neurons (*Aizawa et al., 2001*), levels of p-cofilin rose rapidly above 3-fold in the GC of control cortical neurons during the first minute after exposure to Sema 3A, but underwent a sharp reduction during the next four minutes to 43% of basal levels (*Figure 7D,E*). This sharp F-actin stabilizing period followed by a fast increase in F-actin depolymerization is thought to reflect the reorganization of the actin cytoskeleton during GC collapse (*Aizawa et al., 2001*). Consistent with this plausible signaling cascade and lesser GC collapse due to a diminished response to Sema 3A, GCs of both AnkB440 KO and p.Y1229H L1CAM neurons showed a different pattern of cofilin phosphorylation. First, basal p-cofilin signal per GC area was roughly 40% and 30% lower in AnkB440 KO and p.Y1229H L1CAM neurons, respectively, relative to control (*Figure 7D,E*), which is consistent with lower levels of p-cofilin in AnkB440 brains (*Figure 7A,C*). Second, in contrast to the higher than three-fold increase in control neurons, the rise in p-cofilin during the first minute of Sema 3A treatment was below two-fold in both AnkB440 KO and p.Y1229H L1CAM GCs, which represented only 30-40% of control levels. Lastly, in AnkB440 KO and p.Y1229H L1CAM GCs the reduction of p-cofilin five minutes past Sema 3A treatment was approximately only 20% lower than its peak at one minute and remained around 50% above basal levels (*Figure 7D,E*). These aberrant patterns of cofilin regulation indicate that AnkB440 and its association with L1CAM promote steps of the Sema 3A signal transduction pathways upstream of changes in cofilin activation and F-actin disassembly during GC collapse.

### Autism-linked *ANK2* variants affect the transduction of Sema 3A repulsive cues

The results above support a mechanism wherein AnkB440 is required to modulate the initiation, growth, and establishment of axon collateral branches and to properly respond to Sema 3A repulsive cues that modulate GC collapse, axon guidance, and pruning. Consequently, AnkB440 deficiencies can result in aberrant structural and functional axon connectivity, which in turn may contribute to the pathogenicity of *ANK2* variants in ASD. AnkB440^R2589fs^ mice, which models the *de novo* p.(R2608fs) frameshift variant in exon 37 of *ANK2* found in an individual diagnosed with ASD, exhibit ectopic axonal connections assessed by brain DTI and axonal hyperbranching *in vitro* (*Yang et al., 2019*). Instead of full-length AnkB440, AnkB440^R2589fs^ mice express a 290 kDa truncated product that lacks portions of the NSD unique to AnkB440 plus the entire death and C-terminal regulatory domains (*Yang et al., 2019*) (*Figure 8A,B asterisk, see Figure 8-source data 1, 2, 5*). Given that this truncated protein fails to associate with L1CAM *in vivo* (*Yang et al., 2019*), we tested whether its expression could also disrupt Sema 3A-induced GC collapse. Like AnkB440 KO and p.Y1229H L1CAM neurons, AnkB440^R2589fs^ neurons had diminished GC collapse response to Sema 3A (*Figure 8C,D*). Thus, in addition to altered microtubule stability, impaired responses to Sema 3A during GC collapse likely contribute to the axonal branching and connectivity deficits observed in AnkB440^R2589fs^ brains and neurons.

**Figure 8.**
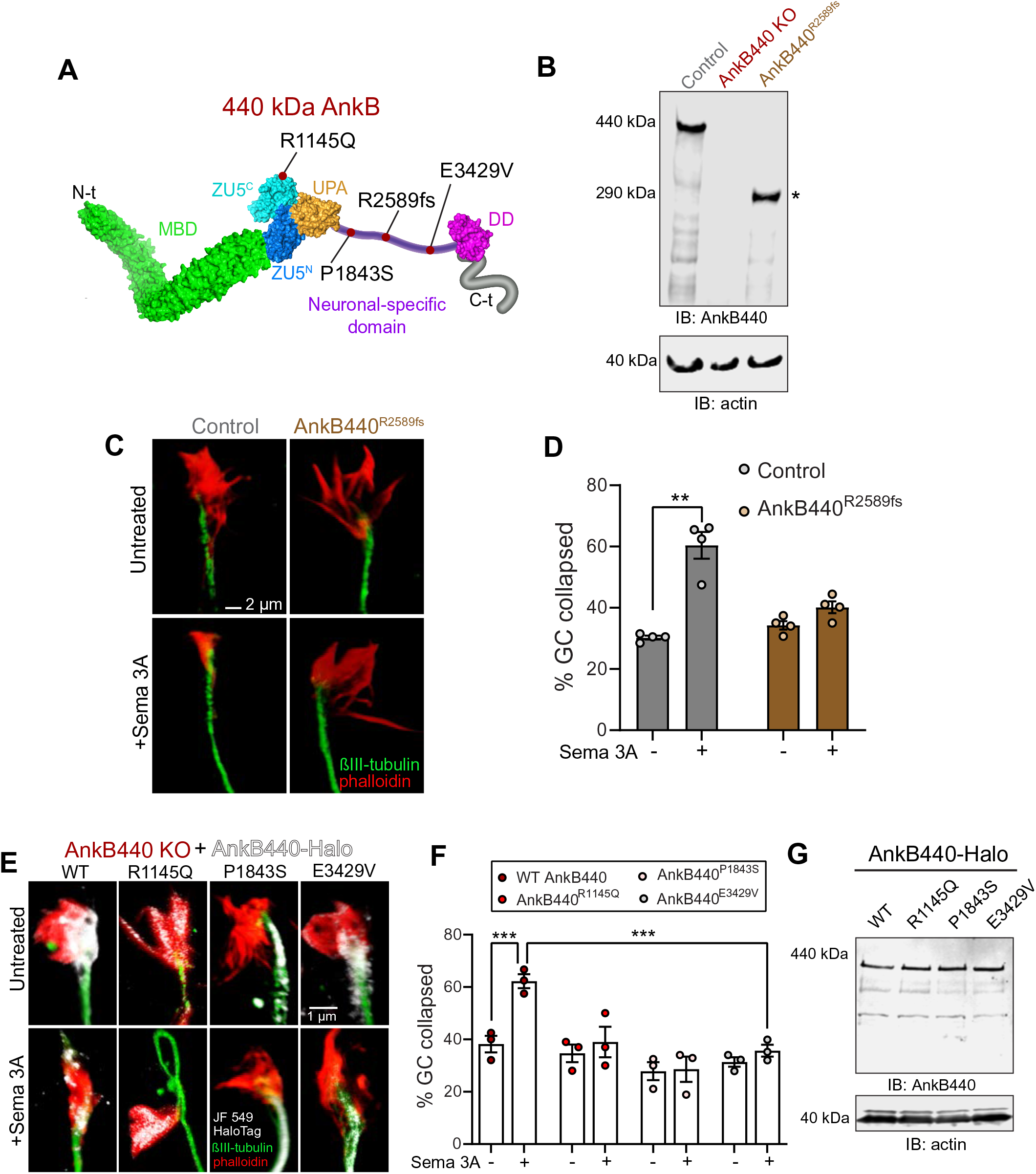
Autism-linked AnkB440 variants affect the transduction of Sema 3A cues during GC collapse. (**A**) Red dots indicate the position within AnkB440 functional domains of the ASD-linked *ANK2* variants evaluated. (**B**) Western blot analysis of expression of AnkB440 in cortical lysates of PND1 control mice of the indicated genotypes assessed with an AnkB440-specific antibody. Brains of AnkB440^R2589fs^ mice express a truncated 290-kDa AnkB440 fragment (asterisk). (**C**) Images of the distal portion of the axon of control and AnkB440^R2589fs^ neurons untreated and treated with Sema 3A and stained with phalloidin and βIII-tubulin. Scale bar, 2 μm. (**D**) Percent of axon GCs that collapse before and after Sema 3A treatment. Data represent mean ± SEM collected from an average of n=130 GCs/treatment condition/genotype from four independent experiments. Unpaired *t* test. **p < 0.01. (**E**) Images of the distal portion of the axon of DIV3 AnkB440 KO cortical neurons rescued with the indicated AnkB440-Halo plasmids untreated and treated with Sema 3A. Scale bar, 1 μm. Percent of axon GCs that collapse before and after Sema 3A treatment. Data represent mean ± SEM collected from an average of n=70 GCs/treatment condition/genotype from three independent experiments. Unpaired *t* test. ***p < 0.001. (F) Western blot analysis of expression of AnkB440-Halo plasmids in HEK293T cells.

Over 70 *ANK2* variants have been identified in individuals diagnosed with ASD (*De Rubeis et al., 2014; Iossifov et al., 2014; Iossifov et al., 2015*). ASD-linked *ANK2* variants target the NSD or domains shared by both AnkB440 and AnkB220 isoforms. We previously reported that *de novo* ASD variants p.(P1843S) and p.(E3429V) in the NSD of AnkB440 (*Figure 8A*) failed to rescue axon hyperbranching of AnkB440 KO cortical neurons (*Yang et al., 2019*). Therefore, we evaluated whether these AnkB440-specific variants could restore GC responses to Sema 3A of AnkB440 KO neurons. We also tested expression of Halo-tagged AnkB440 bearing the *de novo* variant p.(R1145Q) ASD variant in exon 30, which encodes a portion of the ZU5^C^ domain common to both AnkB220 and AnkB440 (*Figure 8A*). AnkB440 KO cortical neurons expressing these ASD-linked AnkB440 variants failed to rescue GC collapse in response to Sema 3A (*Figure 8E,F)*. These variants do not appear to affect protein stability, given that they expressed normal levels of full-length AnkB440 protein when transfected in HEK293T cells, which lack endogenous AnkB440 expression (*Figure 8G, see Figure 8-source data 3-5*). Instead, they likely disrupt AnkB440 distribution, as in the case of the p.(P1843S) AnkB variant, which exhibited reduced Halo-AnkB440 localization to the GCs (*Figure 8E,F*), or might disrupt interaction with specific partners. Together, these results suggest that AnkB440 variants can lead to loss-of-function effects that may affect structural neuronal connectivity and contribute to pathogenicity in ASD.

## Discussion

Variants in ankyrin genes have emerged as risk factors in multiple neurodevelopmental and psychiatric disorders. For example, several GWAS and other genetic studies have found both rare and common variants in *ANK3*, which encodes ankyrin-G (AnkG), associated with bipolar disorder (*Baum et al., 2008; Ferreira et al., 2008*) and schizophrenia (*Cruz et al., 2009; Schizophrenia Psychiatric Genome-Wide Association Study (GWAS) Consortium, 2011*). Similarly, variants in *ANK2* have been identified in individuals with ASD and intellectual disability (*De Rubeis et al., 2014; Iossifov et al., 2014; Iossifov et al., 2015*) and it is ranked as a top high confidence ASD gene with one of the highest mutability scores (*Ruzzo et al., 2019*). Despite their structural similarities and degree of sequence conservation, ankyrins diverge in cell type expression, subcellular localization, and protein partners in the brain (*Lorenzo, 2020*). For instance, AnkG preferentially localizes to the axon initial segment (AIS), where it acts as the master regulator of AIS organization, while AnkB is widely distributed throughout the axon (*Lorenzo et al., 2014; Yang et al., 2019; Lorenzo, 2020*). Ankyrins achieve a second order of functional specialization through alternative splicing, which yields giant isoforms with unique inserted sequences in both AnkG and AnkB in neurons (*Bennet and Lorenzo, 2016*; *Lorenzo, 2020*). ASD-linked AnkB variants fall both within the inserted region unique to AnkB440 and in domains shared by AnkB440 and AnkB220 isoforms.

The present study sheds new light into the isoform-specific functions of AnkB in modulating axonal architecture and guidance in the developing brain and the potential contribution of AnkB deficits to ASD pathology. We previously showed that expression of the truncated product of the frameshift mutation p.P2589fs that models the *de novo* p.(R2608fs) ASD variant in AnkB440 causes stochastic increases in structural cortical connectivity in mouse brains and exuberant collateral axon branching in cortical neuron cultures (*Yang et al., 2019*). We confirmed the development of axon hyperbranching in cultured neurons selectively lacking AnkB440 (*Chan et al., 2020*), but not in neurons from AnkB220 KO mice. Using Golgi staining, we determined that AnkB440 KO, but not AnkB220 KO, cortical neurons grow more collateral branches, consistent with the *in vitro* results. We also found thickening of the corpus callosum in AnkB440 KO mice, which could result from volumetric gains caused by hyperbranching of callosal axons. These findings support critical and specialized roles of AnkB440 in modulating axon collateral branch formation and pruning.

Previous results implicated AnkB440 in suppressing collateral branch formation via the stabilization of microtubule bundles near the plasma membrane (*Yang et al., 2019; Chan et al., 2020*). Loss of AnkB440 or expression of the 290-kDa truncated AnkB440 product promotes microtubule unbundling, which facilitates microtubule invasion of the nascent filopodia, a necessary step in the commitment to forming a new branch (*Yu et al., 1994*). The formation of transient actin nucleation sites marks the point of formation and precedes the emergence of the precursor filopodia (*Gallo, 2011*). In this study, we show that selective loss of AnkB440 KO in cultured cortical neurons results in larger number of actin patches relative to control and AnkB220 KO neurons, which correlated with higher number of axonal filopodia and collateral branches. Thus, AnkB440 modulates collateral branch formation through a combined mechanism that suppresses the stochastic formation of collateral filopodia by lowering the density of F-actin-rich branch initiation points, as well as the invasion of microtubules into the maturing filopodia. Although F-actin patches serving as precursor to filopodia have been observed *in vitro* (*Loudon et al., 2006; Spillane et al., 2011*) and *in vivo* (*Spillane et al., 2011; Hand et al., 2015*), little is known about the factors that regulate their formation and dynamics (*Armijo-Weingart and Gallo, 2017*). How AnkB440 modulate actin patches is not clear. The crosstalk between actin and microtubule networks have been proposed to orchestrate the emergence of branches (*Dent and Kalil, 2001*; *Pacheco and Gallo, 2016*), and splaying of microtubule bundles have been observed to correlate with F-actin accumulation at branch points (*Dent and Kalil, 2001*). Thus, it is possible that microtubule unbundling resulting from AnkB440 deficits promotes the seeding and growth of F-actin patches at sites of axon branch formation. Alternatively, AnkB440 may associate with actin through their common partner βII-spectrin, although loss of βII-spectrin does not lead to axonal hyperbranching *in vitro* and may not affect the number or dynamics of actin patches (*Lorenzo, et al., 2019*). While loss of AnkB220 does not affect axonal branching *in vitro* and in mouse brains, its expression in AnkB440 null neurons may be necessary to sustain hyperbranching. This is possibly due to the role of AnkB220 in axonal organelle and vesicle transport and growth, which are not affected by exclusive loss of AnkB440.

While cell-autonomous factors that modulate axonal branching *in vitro*, such as the formation of actin patches, correlate with the number of axon branches (*Gallo, 2011*), live imaging of actin dynamics during *in vivo* axonal development found that that correlation does not hold in projection neurons in the mouse cortex (*Hand et al., 2015*). Instead, the number of collateral axon branches along the axon is determined by the cortical layer they transverse, indicating that extrinsic factors that modulate branch formation and pruning may be involved in determining the pattern of axonal innervation (*Hand et al., 2015*). Several reports support the activity of Sema 3A as an extrinsic repellent cue that inhibits axon branching *in vitro* (*Dent et al., 2004*) and prunes cortical axons and collateral axon branches *in vivo* (*Polleux et al., 1998; Bagri et al., 2003*). We show that AnkB440 is required to transduce repellent Sema 3A signals *in vitro* to facilitate the collapse of GCs from the axon and collateral branches, which offers a plausible cellular mechanism underlying the ectopic neuronal connectivity and axonal hyperbranching observed in AnkB440-deficient mouse brains. AnkB440-mediated transduction of Sema 3A during GC collapse involves AnkB440 binding to L1CAM, which in turn stabilizes the Sema 3A holoreceptor complex composed of L1CAM-Nrp1 at the cell surface (*Castellani et al., 2000; Castellani et al., 2002; Castellani et al., 2004*). Interestingly, GCs of neurons lacking the F-actin and AnkB partner βII-spectrin respond normally to Sema 3A, which indicates that βII-spectrin is not required for Sema 3A-induced GC collapse. However, given βII-spectrin’s role in the development and wiring of axons in mouse brains (*Galiano et al., 2012; Lorenzo et al., 2019*) and the recent identification that pathogenic variants in *SPTBN1*, which encodes βII-spectrin, cause a neurodevelopmental syndrome associated with deficits in cortical connectivity (*Cousin et al., 2020*), we cannot rule out its involvement in axonal guidance through alternative mechanisms. Interestingly, although AnkB440 binding to microtubules suppresses branch initiation, loss of this interaction does not affect the response to Sema 3A during GC collapse. Instead, transduction of Sema 3A signaling via the AnkB440-L1CAM-Nrp1 complex modulates F-actin dynamics through LIMK phosphorylation of cofilin.

Our structure-function studies in AnkB440 KO cortical neurons found that *de nov*o ASD variants p.(R1145Q), p.(P1843S), p.R2589fs, and p.(E3429V) in AnkB440 fail to restore cellular responses to Sema 3A during GC collapse. Interestingly, expression of the p.R2589fs variants caused axon hyperbranching *in vitro* and ectopic axon connectivity *in viv*o, and the remaining variants did not rescue normal collateral branching in AnkB440 KO neurons *in vitro* (*Yang et al., 2019*). The 290-KDa AnkB440 product of p.R2589fs is expressed at slightly reduced levels compared to full-length AnkB440 and shows reduced binding to L1CAM *in vivo* (assessed through PLA) despite preserving the L1CAM binding region in the membrane binding domain (MBD) (*Yang et al., 2019*). This diminished AnkB440^R2589fs^-L1CAM association may result from the lower expression of the truncated AnkB440 protein (*Yang et al., 2019*), although it is also possible that this truncation may induce changes in the conformational or self-regulation of truncated AnkB440 that might interfere with its association with L1CAM. The mechanisms by which the other ASD AnkB440 variants evaluated disrupt AnkB440 function are less clear. The p.(P1843S) might alter AnkB440 distribution. This variant also falls within the microtubule binding region of AnkB440 (*Chen et al., 2020*) and may affect microtubule binding, which if true, may explain the axon hyperbranching phenotype, but it is not clear how it interferes with GC collapse. Both ASD variants p.(R1145Q) and p.(E3429V) in AnkB440 reside outside of the MBD and the microtubule-binding regions. p.(R1145Q) is located in the ZU5^C^ domain shared by AnkB220 and AnkB440 and it may affect AnkB’s ability to bind PI3P lipids, given its proximity to the PI3P lipids binding region (*Lorenzo et al., 2014*). However, while AnkB220 requires binding to PI3P lipids to associate with intracellular membranes and enable axonal transport (*Lorenzo et al., 2014*), AnkB440 does not modulate axonal transport. In future experiments it would be important to determine whether AnkB440 binds PI3P lipids and the significance of AnkB440-PI3P lipid binding activity for axonal development and axonal connectivity in the developing brain.

Besides the significance of *ANK2* in ASD, neuronal pathways involving the AnkB440-L1CAM complex are also relevant to other neurological diseases. For example, a few hundred variants in L1CAM have been described in individuals with CC hypoplasia, retardation, adducted thumbs, apasticity and hydrocephalus (CRASH) syndrome (*Rosenthal et al., 1992; Jouet et al., 1994; Weller and Gärtner, 2001; Vos et al., 2010*). Pathological L1CAM variants can affect both its extra- and intracellular domains and disrupt binding to molecular partners including AnkB and Nrp1 (*Schäfer and Altevogt, 2010*). Studies in mouse models that constitutively lack L1CAM have reported CC hypoplasia, cerebellar and other brain malformations, and axon guidance defects in the corticospinal tract (*Dahme et al., 1997; Cohen et al., 1998; Fransen et al., 1998*). Interestingly, neurons differentiated from human embryonic stem (ES) cells in which expression of endogenous LICAM was knockdown through homologous recombination showed reduced axonal length and deficient axonal branching relative to matching control neurons (*Patzke et al., 2016*). These results are in contrast with the cellular phenotypes we observe in AnkB440 KO neurons, even though ES cell-derived L1CAM KO neurons showed noticeable downregulation of AnkB, which appear to have been largely driven by significant loss of AnkB440. We found that loss of AnkB440 reduces L1CAM abundance at the cell surface but does not significantly change total levels of L1CAM in the cortex and in cortical neuron cultures. L1CAM expression is also normal in brains of AnkB440^R2589fs^ mice (*Yang et al., 2019*). While it is plausible that normal AnkB expression requires L1CAM, and not the reverse, it would be important to confirm whether these changes in protein expression are specific to human ES cell-derived neurons. The phenotypic differences between AnkB440- and L1CAM-deficient mice point to additional, independent pathways involving AnkB440 and L1CAM, including the functional relationship of L1CAM with other ankyrins, which collectively underscore the importance of these proteins in neuronal structure and signaling. For instance, a recent report implicates an L1CAM-AnkG association at the AIS of neocortical pyramidal neurons in their innervation by GABAergic chandelier cells (*Tai et al., 2019*). AnkB440 it is widely distributed through all axonal domains, including the AIS. Further work will be needed to determine whether AnkB440-L1CAM complexes localize at the AIS and if they contribute to this specific type of pyramidal neuron innervation.

In summary, our findings unveil a critical role of AnkB440 and its binding to L1CAM in promoting the clustering of L1CAM-Nrp1 complexes at the cell surface of GCs of cortical neurons. Accordingly, we show that the AnkB440-L1CAM-Nrp1 complex enables the chemorepellent action of Sema 3A to induce GC collapse in axons and collateral branches, thereby providing novel insight into the mechanisms of axonal guidance and branch pruning. As we show, *ANK2* variants cause disruption of this signaling axis, which might lead to cortical miswiring and contribute to the neuropathology of ASD.

## Methods

### Mouse lines and animal care

Experiments were performed in accordance with the guidelines for animal care of the Institutional Animal Care and Use Committee at the University of North Carolina at Chapel Hill. The total AnkB knockout mice (total AnkB KO), the conditional AnkB440 (AnkB440^flox^) mouse line, the knock-in AnkB440^R2589fs^ mouse line that models a human ASD-linked *ANK2* variant, and the mice carrying the Y1229H mutation in L1CAM were a gift from Dr. Vann Bennett at Duke University and have been previously reported (*Scotland et al., 1998; Lorenzo et al., 2014; Yang et al., 2019; Chan et al., 2020*). βII-spectrin floxed mice (*Sptbn1^flox/flox^*, a gift from Dr. Mathew Rasband) have been previously reported (*Galiano et al., 2012; Lorenzo et al., 2019*). AnkB440 KO and AnkB220 KO (described below) mice respectively lacking AnkB440 and AnkB220 in neural progenitors were generated by crossing *AnkB440^flox/flox^* or *AnkB220^flox/flox^* animals to the Nestin-Cre line [B6.Cg-Tg(Nes-cre)1Kln/J, stock number 003771] from The Jackson Laboratory. A similar breeding strategy was used to generate mice with loss of βII-spectrin in neural progenitor. All mice were housed at 22°C ± 2°C on a 12-hour-light/12-hour-dark cycle and fed ad libitum regular chow and water.

### Generation of conditional AnkB220 knockout mice

Mice carrying a floxed allele that selectively targets the AnkB220 isoform (*AnkB220^flox/flox^*) were generated by the Animal Model Core at the University of North Carolina at Chapel Hill using CRISPR/Cas9-mediated integration of a targeting vector into mouse embryonic stem (ES) cells, followed by ES cell injection into blastocytes and production of chimeric progeny. In brief, the CCTop website (https://crispr.cos.uni-heidelberg.de) was used to identify potential Cas9 guide RNAs targeting *Ank2* intron 35 and the 5’ end of exon 37. Selected guide RNAs were cloned into a T7 promoter vector followed by *in vitro* transcription and spin column purification. Functional testing was performed by transfecting Cas9 protein/guide RNA ribonucleoprotein complexes into a mouse embryonic fibroblast cell line. The guide RNA target regions were amplified from transfected cells and analyzed by T7endo1 assay (NEB) to detect genome editing activity at the target site. Guide RNAs selected for genome editing in mouse embryonic stem cells were *Ank2*-i35-sg73T (protospacer sequence 5’-GGTTCTAGTCTTCCCGA-3’) and *Ank2*-E37-sg79B (protospacer sequence 5’-GTCCGGACTTGCTAAGAC-3’). A donor vector was constructed for homologous recombination that included a 1002 bp 5’ homology arm corresponding to the sequence immediately 5’ of the cut site of Cas9/*Ank2*-i35-sg73T; a LoxP site; 368 bp 3’ end of *Ank2* intron 35 including splice acceptor sequence; 3707 bp cDNA encompassing exons 36 and 38-46 (isoform lacking exon 37); a 814 bp stop cassette comprised of 3 tandem copies of SV40 polyadenylation sequence; a FRT-flanked selection cassette with PGK mammalian promoter, a EM7 bacterial promoter, a neomycin resistance gene and PGK polyadenylation cassette, all in reverse orientation relative to the *Ank2* elements; a LoxP site; a second 368 bp 3’ end of Ank2 intron 35 including splice acceptor sequence; a 86 bp segment including 22 bp exon 36 fused to 64 bp 5’ end of exon 37; silent point mutations designed to disrupt the Ank2-E37-sg79B target site. The sequence GTCTTA was mutated to GTGCTC, corresponding to mutation of a valine codon from GTC to GTG and a leucine codon from TTA to CTC; and a 996 bp 3’ homology arm corresponding to sequences immediately 3’ of the silent point mutations.

The donor vector was incorporated into C57BL/6N ES cells by nucleofection with 3 µM Cas9 protein (Thermo Scientific), 1.6 25 µM each Ank2-i35-sg73T and Ank2-E37-sg79B guide RNAs and 200 ng/µl (20 µg total) circular donor vector DNA. Cells were selected on G418 and resistant clones were screened for homologous integration of the donor vector at the *Ank2* locus. PCR-positive clones were analyzed by Southern blot with 5’ and 3’ external probes and neomycin cassette internal probe. Two clones, F6 and H10, were identified with homologous integration of the donor. Targeted ES cell clones F6 and H10 were injected in Albino C57BL/6N blastocysts for chimera production. Chimeras were mated to transgenic animals expressing Flp recombinase on Albino C57BL/6N genetic background. Germline transmission of the targeted allele was obtained from both clones, although clone H10 gave more germline transmission pups. Clone H10 had an apparent random integration event in addition to the homologous event. Therefore, pups were screened to identify clones with the homologous integration event in absence of the random integration event. Selected founders were bred for five generations to C57BL/6J mice after which heterozygous carriers of the *Ank2* targeting allele (*AnkB220^flox/+^*) were bred to each other to generate homozygous carriers (*AnkB220^flox/flox^*). The WT *Ank2* allele was identified by PCR using primers ABCS-WT-F (5’-GCTTTGTTGTATGTATGAATGTGCTAC-3’) and ABCS-WT-R (5’-TTCCTCATCGCTGACAATAACC-3’), which produce a 328 bp DNA fragment. The *AnkB220^flox^* allele was detected by PCR using primers ABCS-RE-F2 (5’-GCTTGGCTGTGTTCACAAACA-3’) and ABCS-RE-R2 (5’-GACTTGCGAGCACAGGAACTT-3’), which produce a 639 bp DNA fragment.

### Plasmids

Plasmids used for transfections include pmCherry-C1 (Clontech), pLAMP1-mGFP (Addgene #34831, gift from Dr. Esteban Dell’Angelica), pmCherry-Lifeact-7 (Addgene #54491, gift from Dr. Michael Davidson) and pcDNA3.1-L1CAM (Addgene # 12307, gift from Dr. Vance Lemmon). Plasmids pBa-AnkB440-Halo, pBa-AnkB440-PSK-Halo, pBa-AnkB440^P1843S^-Halo, pBa-AnkB440^E3429V^ (gifts from Dr. Vann Bennett) have been previously described (*Yang et al., 2019; Chan et al., 2020*). The pBa-AnkB440^R1145Q^-Halo plasmid was generated via GeneArt™ site-directed mutagenesis (Life Technologies) using primers R1145Q_F (5’-CGCATCATCACCCAAGACTTCCCACAG-3’) and R1145Q_R (5’-CTGTGGGAAGTCTTGGGTGAT GAT GCG-3’). To generate pCAG-AnkB220-GFP, an XhoI and NotI fragment containing the AnkB220 cDNA was cloned into the same sites of pCAG-eGFP-N1. The GFP-tagged AnkB440 vector was generated by digesting pBa-AnkB440-Halo with NheI and MfeI to excise the C terminal Halo tag. GFP was amplified with complementary ends from pCAG-AnkB220-GFP using high fidelity Phusion polymerase (Takara) and primers F-NheI: 5’-CAACAATGAGGCTAGCCGGGATCCACCGGTCGCC-3’ and R-MfeI: 5’-TTAACAACAACAATTGATCTAGAGTCGCGGCCGC-3’). Linearized fragments were assembled using In-Fusion Cloning reagents (Takara). All plasmids were verified by full-length sequencing prior to transfection.

### Antibodies and fluorescent dyes

Affinity-purified rabbit pan anti-AnkB and anti-AnK440 antibodies used at a 1:500 dilution for immunohistochemistry and 1:5000 for western blot, were generated by Dr. Vann Bennett laboratory and have been previously described (*Lorenzo et al., 2014; Yang et al., 2019; Chan et al., 2020*). Other antibodies used for western blot analysis included rabbit anti-Sema 3A (1:500, #ab23393) and rabbit anti-Nrp1 (1:1,000, #ab205718) all from Abcam. We also used rabbit anti-Plexin A1 (1:200, # APR-081, Alomone), rabbit anti-phospho-Cofilin (Ser3) (1:1,000, #3311) and rabbit anti-LIMK1 (1:1,000, #3842) from Cell Signaling, mouse anti-cofilin (1:1,000, clone 1G6A2, Proteintech), mouse pan anti-actin (1:2,000, clone C4, #MAB1501, Millipore-Sigma), and mouse anti-GluR1 (1:500, clone N355/1, #75-327, NeuroMab). Commercial antibodies used for immunofluorescence included mouse anti-neurofilament (1:200, clone SMI-312, #837904) and mouse anti-βIII-tubulin (1:100, clone TU-20, #MAB1637) from Millipore-Sigma, chicken anti-GFP (1:1000, #GFP-1020) from Aves, and mouse anti-Satb2 (1:200, clone SATBA4B10, # ab51502), rat anti-Ctip2 (1:500, clone 25B6, # ab18465) and rabbit anti-Tbr1 (1:200, # ab31940), all from Abcam. The mouse anti-L1CAM (clone 2C2, # ab24345) was used at 1:200 for both western blot and immunofluorescence analyses. Detection of surface Nrp1 by immunofluorescence was conducted using rabbit anti-Nrp1 extracellular antibody (1:50, # ANR-063, Alomone) that recognizes amino acid residues 502-514 in the extracellular N-terminus of Nrp1. The presence of the Halo tag was detected by incubation with the JF 549 HaloTag ligand from Janelia Farm. F-actin was labeled with phalloidin conjugated to Alexa Fluo-647, 568, and 488 dyes (Life Technologies). Secondary antibodies purchased from Life Technologies were used at 1:400 dilution for fluorescence-based detection by confocal microscopy. Secondary antibodies included donkey anti-rabbit IgG conjugated to Alexa Fluor 568 (#A10042), donkey anti-mouse IgG conjugated to Alexa Fluor 488 (#A21202), goat anti-chicken IgG conjugated to Alexa Fluor 488 (#A11039), donkey anti-rat IgG conjugated to Alexa Fluor 647 (#A21247), goat anti-rat IgG conjugated to Alexa Fluor 568 (#A11077), donkey anti-mouse IgG conjugated to Alexa Fluor 568 (#A10037), donkey anti-rabbit IgG conjugated to Alexa Fluor 647 (#A31573), goat anti-rabbit IgG conjugated to Alexa Fluor 594 (#R37117), and goat anti-mouse IgG conjugated to Alexa Fluor 488 (#A11001).Fluorescent signals in western blot analysis were detected using goat anti-rabbit 800CW (1:15000, #926-32211) and goat anti-mouse 680RD (1:15000, #926-68070) from LiCOR.

### Plasmid transfection for biochemistry analysis

Transfection of Halo-tagged AnkB440 plasmids were conducted in HEK293T cells grown in 10 cm culture plates using the calcium phosphate transfection kit (Takara) and 8 µg of plasmid. Cell pellets were collected 48 hours after transfection.

### Inmunoblots

Protein homogenates from mouse brains or transfected cells were prepared in 1:9 (wt/vol) ratio of homogenization buffer (8M urea, 5% SDS (wt/vol), 50mM Tris pH 7.4, 5mM EDTA, 5mM N-ethylmeimide, protease and phosphatase inhibitors) and heated at 65°C for 15 min to produce a clear homogenate. Total protein lysates were mixed at a 1:1 ratio with 5x PAGE buffer (5% SDS (wt/vol), 25% sucrose (wt/vol), 50mM Tris pH 8, 5mM EDTA, bromophenol blue) and heated for 15 min at 65°C. Samples were resolved by SDS-PAGE on 3.5-17.5% acrylamide gradient gels in Fairbanks Running Buffer (40mM Tris pH 7.4, 20mM NaAc, 2mM EDTA, 0.2% SDS (wt/vol)). Proteins were transferred at 29V overnight onto 0.45 μm nitrocellulose membranes (#1620115, BioRad) at 4°C in methanol transfer buffer (25mM Tris, 1.92M Glycine, 20% methanol). Transfer efficiency was determined by Ponceau-S stain. Membranes were blocked in TBS containing 5% non-fat milk for 1 hour at room temperature and incubated overnight with primary antibodies diluted in antibody buffer (TBS, 5% BSA, 0.1% Tween-20). After 3 washes in TBST (TBS, 0.1% Tween-20), membranes were incubated with secondary antibodies diluted in antibody buffer for two hours at room temperature. Membranes were washed 3x for 10 minutes with TBST and 2x for 5 minutes in TBS. Protein-antibody complexes were detected using the Odyssey® CLx Imaging system (LI-COR).

### Labeling and detection of biotinylated surface proteins

Cortical neuronal cultures were washed three times with ice-cold PBSCM (PBS + 1mM MgCl_2_ + 0.1 mM CaCl_2_) and incubated with 0.5 mg/ml Sulfo-NHS-SS-biotin (Life Technologies) for 1 hour at 4°C. Reactive biotin was quenched by two consecutive 7-minute incubations with 20 mM glycine in PBSCM on ice. Cell lysates were prepared in TBS containing 150 mM NaCl, 0.32 M sucrose, 2 mM EDTA, 1% Triton X-100, 0.5% NP40, 0.1% SDS, and complete protease inhibitor cocktail (Sigma). Cell lysates were incubated with rotation for 1 hour at 4°C and centrifuged at 100,000 x g for 30 min. Soluble fractions were collected and incubated with high capacity NeutrAvidin™ agarose beads (Pierce) overnight at 4°C to capture biotinylated surface proteins. Beads were washed three times with TBST. Proteins were eluted in 5x-PAGE buffer and resolved by SDS-PAGE and western blot.

### Histology and immunohistochemistry

Brains were fixed by transcardial perfusion with 4% PFA in PBS before overnight immersion in the same fixative at 4°C. After fixation, brains were stored in PBS at 4°C until use. Brains were then transferred to 70% ethanol for 24 hours and paraffin embedded. 10 µm coronal brain sections were cut using a microtome (Leica RM2135) and mounted on glass slides. Sections were deparaffinized and rehydrated using a standard protocol of washes: 3 x 3 xylene washes, 3 x 2 min 100% ethanol washes, and 1 x 2 min 95%, 80%, 70%, ethanol each, followed by ≥5 min in PBS. Sections were processed for antigen retrieval using 10 mM sodium citrate with 0.5% Tween-20, pH 6 in a pressure cooker for 3 minutes at maximum pressure. Sections were cooled, washed in PBS and blocked in antibody buffer (4% BSA, 0.1% Tween-20 in PBS) for 90 minutes at room temperature. Tissue sections were then incubated with primary antibody in antibody buffer overnight at 4°C and with secondary antibodies for 2 hours at room temperature, washed with PBS, incubated with DAPI where applicable, and mounted with Prolong Gold Antifade Reagent (Life Technologies).

### Golgi stain

Golgi staining of PND25 brains was conducted using the FD Rapid GolgiStain Kit (FD Neurotechnologies Inc.) In brief, brains were immersed in Solutions A+B for 2-3 weeks, before being transferred to solution C for 3-6 days. 100 µm coronal cryosections from areas of the somatosensory cortex were collected on gelatin-coated microscope slides, counter-stained following manufacturer recommendations, and mounted in Permount for imaging.

### Primary neuron culture

Primary cortical neuronal cultures were established from E16-PND0 mice. Cortices were dissected in Hibernate E (Life Technologies) and digested with 0.25% trypsin in HBSS (Life Technologies) for 20 min at 37°C. Tissue was washed three times with HBSS, dissociated in DMEM (Life Technologies) supplemented with 5% fetal bovine serum (FBS, Genesee), and gently triturated through a glass pipette with a fire-polished tip. Dissociated cells were filtered through a 70 µm cell strainer to remove any residual non-dissociated tissue and plated onto poly-D-lysine-coated 1.5 mm coverglasses or dishes (MatTek) at a density of 4×10^4^ cells/cm^2^ for transfection and imaging experiments. For all cultures, media was replaced 3 hours after plating with serum-free Neurobasal-A medium containing B27 supplement (Life Technologies), 2 mM Glutamax (Life Technologies), and 1% penicillin/streptomycin (Life Technologies) (neuronal growth media). 5μM cytosine-D-arabinofuranoside (Sigma) was added to the culture medium to inhibit the growth of glial cells three days after plating. Neurons were maintained at 37°C with 5% CO_2_ and fed twice a week with freshly made culture medium until use.

### Plasmid transfection for time-lapse live imaging and immunofluorescence analysis

For neuronal rescue experiments, AnkB440 KO primary cortical neurons were transfected with Halo-tagged AnkB440 or GFP-tagged AnkB220 plasmids at DIV0 by lipofection (Lipofectamine 2000, Thermo Fisher Scientific). After brain dissociation, 2.5×10^5^ cells/ml neurons were incubated with Lipofectamine 2000 (12 µl/ml) and plasmid DNA (2.5 µg/ml) in suspension at 37°C for 45 min. Cells were pelleted at 200 g for 4 minutes and plated in poly-D-lysine-coated coverglasses or dishes and used for studies at DIV3. For time-lapse imaging experiments DIV5 cortical neurons were co-transfected with 1µg of pLAMP1-GFP using lipofectamine 2000 and imaged 48-96 hours after transfection. For experiments that evaluate axonal length and branching, DIV0 or DIV2 neurons were transfected with 500 ng of pmCherry-C1. Neurons were processed for immunofluorescence 3 days after transfection. Evaluation of L1CAM binding to GFP-tagged AnkB220 or AnkB440 through a membrane recruitment assay was conducted HEK293T in cells independently transfected with 100 ng of pcDNA3.1-L1CAM, GFP-AnkB220, or GFP-AnkB440 plasmids, or co-transfected with 100 ng of pcDNA3.1-L1CAM in combination with either GFP-AnkB220 or GFP-AnkB440 plasmids 48 hours post-transfection.

### Growth cone collapse experiments

DIV3 neurons were treated with Sema 3A (250 ng/ml, Peprotech, Cat. 150-17H) or Ephrin A5 (1 µg/ml, R&D Systems, Cat. 374-EA) for 1, 5, 15, or 30 min in neuronal growth media, fixed for 15 min with 4% PFA/4% sucrose, and washed in PBS. Neurons were permeabilized with 0.2% Triton X-100 in PBS for 10 min and blocked in antibody buffer (4% BSA, 0.1% Tween-20 in PBS) for 60 min at room temperature. Neurons were stained with primary antibodies overnight at 4 °C and with a mix of secondary antibodies and phalloidin for two hours at room temperature. Neurons transfected with plasmids expressing Halo-tagged AnkB440 proteins were treated with fresh media containing Sema 3A and with the JF 549 HaloTag ligand (1:200) for 30 min. The percent of collapsed growth cones was recorded for each experimental condition, selecting for transfection when applicable. Axon growth cones were recorded as collapsed based on the presence of a pencil-like shape devoid of lamellae and with a maximum of one filopodia, or intact based on the fan-shaped morphology enriched in filopodia and/or lamellipodia.

### Immunocytochemistry

Neuronal cultures and HEK293T cells were washed with cold PBS, fixed with 4% PFA/4% sucrose for 15 min, and permeabilized with 0.2% Triton-X100 in PBS for 10 min at room temperature. Cells were blocked in antibody buffer for one hour at room temperature and processed for fluorescent staining as tissue sections. For F-actin labeling, Alexa Fluor 488-, Alexa Fluor 568-, or Alexa Fluor 633-conjugated phalloidin (1:100) was added to the secondary antibody mix. To label surface Nrp1, fixed but non-permeabilized DIV3 cortical neurons, untreated and treated with Sema 3A, were blocked as described above and incubated overnight at 44°C with a rabbit anti-Nrp1 antibody that recognizes an extracellular epitope, followed by overnight incubation with donkey anti-rabbit IgG conjugated to Alexa Fluor 568. Neurons were fixed again for 10 min with 4% PFA in PBS, permeabilized for 8 minutes with 0.2% Triton X-100 in PBS, reblocked with antibody buffer for 30 minutes at room temperature and incubated with primary antibodies overnight at 4°C. Secondary antibodies and phalloidin were applied in antibody buffer overnight at °4C. Cells were mounted for imaging with Prolong Gold Antifade Reagent (Life technologies).

### Proximity ligation assay (PLA)

PLA was performed using the commercial Duolink kit (Sigma-Aldrich) following the manufacturer’s recommendations. Fixed and permeabilized neurons were incubated overnight with a pair of primary antibodies specific for the putative interacting partners, each produced in different species. Duolink minus- and plus-probes were used to detect antibody-labeled proteins. In the case of the detection of PLA signal between surface Nrp1 and L1CAM, fixed, but non-permeabilized neurons were first incubated overnight at 4 °C with a rabbit anti-Nrp1 antibody that recognizes an extracellular epitope, followed by cell permeabilization and overnight incubation with mouse anti-L1CAM.

### Image acquisition and image analysis

Brain sections stained with antibodies were imaged using a Zeiss 780 laser scanning confocal microscope (Zeiss) and 405-, 488-, 561-, and 633-nm lasers. Single images and Z-stacks with optical sections of 1 μm intervals and tile scans were collected using the 20x (0.8 NA) and Plan Apochromat 40x oil (1.3 NA) and 63x oil (1.4 NA) objective lenses. Images were processed, and measurements taken and analyzed using NIH ImageJ software. Three-dimensional rendering of confocal Z-stacks was performed using Imaris (Bitplane). Golgi-stained brains were imaged using a Nikon Ti2 Eclipse scope running NIS-Elements. Widefield Z-stacks were taken with 2 µm optical section in the primary somatosensory cortex with a 40x/NA objective.

### Time-lapse video microscopy and movie analyses

Live microscopy of neuronal cultures was carried out using a Zeiss 780 laser scanning confocal microscope (Zeiss) equipped with a GaAsP detector and a temperature- and CO_2_-controlled incubation chamber as previously reported (*Snouwaert et al., 2018*). Movies were taken in the distal axon and captured at a rate of 1 frame/second for time intervals ranging from 60-300 seconds with a 40x oil objective (1.4NA) using the zoom and definite focus functions. Movies were processed and analyzed using ImageJ (http://rsb.info.nih.gov/ij). Kymographs were obtained using the KymoToolBox plugin for ImageJ (https://github.com/fabricecordelieres/IJ_KymoToolBox). In details, space (x axis in µm) and time (y axis in sec) calibrated kymographs were generated from video files. In addition, the KymoToolBox plugin was used to manually follow a subset of particles from each kymograph and report the tracked particles on the original kymograph and video files using a color code for movement directionality (red for anterograde, green for retrograde and blue for stationary particles). Quantitative analyses were performed manually by following the trajectories of individual particles to calculate dynamic parameters including, net and directional velocities and net and directional run length, as well as time of pause or movement in a direction of transport. Anterograde and retrograde motile vesicles were defined as particles showing a net displacement >3 μm in one direction. Stationary vesicles were defined as particles with a net displacement <2 μm.

### Statistical analysis

Sample size (n) for evaluations of growth cone collapse was estimated using power analyses and expected effect sizes based on preliminary data in which we used similar methodologies, specimens, and reagents. We assumed a moderate effect size (f=0.25-0.4), an error probability of 0.05, and sufficient power (1-β=0.8). GraphPad Prism (GraphPad Software) was used for statistical analysis. Two groups of measurements were compared by unpaired, two tailed students *t*-test. Multiple groups were compared by one-way ANOVA followed by Tukey or Dunnett’s multiple comparisons tests.

## Acknowledgments

We thank Dr. Vann Bennett for the gift of the total AnkB KO, AnkB440 KO and L1CAM^Y1229H^ mice and Dr. Matthew Rasband for the gift of the βII-spectrin KO mice. D.N.L. was supported by the University of North Carolina at Chapel Hill School of Medicine as a Simmons Scholar and by the US National Institutes of Health (NIH) grant R01NS110810. Microscopy was performed at the Neuroscience Microscopy Core Facility, supported, in part, by funding from the NIH-NINDS Neuroscience Center Grant P30 NS045892 and the NIH-NICHD Intellectual and Developmental Disabilities Research Center Support Grant U54 HD079124.

## Author contributions

Blake A. Creighton, Formal analysis, Investigation, Methodology, Writing - review and editing; Deepa Ajit, Formal analysis, Investigation, Writing - review and editing; Simone Afriyie, Investigation; Julia Bay, Investigation; Damaris N. Lorenzo, Conceptualization, Data curation, Formal analysis, Investigation, Methodology, Supervision, Funding acquisition, Project administration, Writing - original draft, Writing - review and editing.

---

**Figure 1—figure supplement 1.**
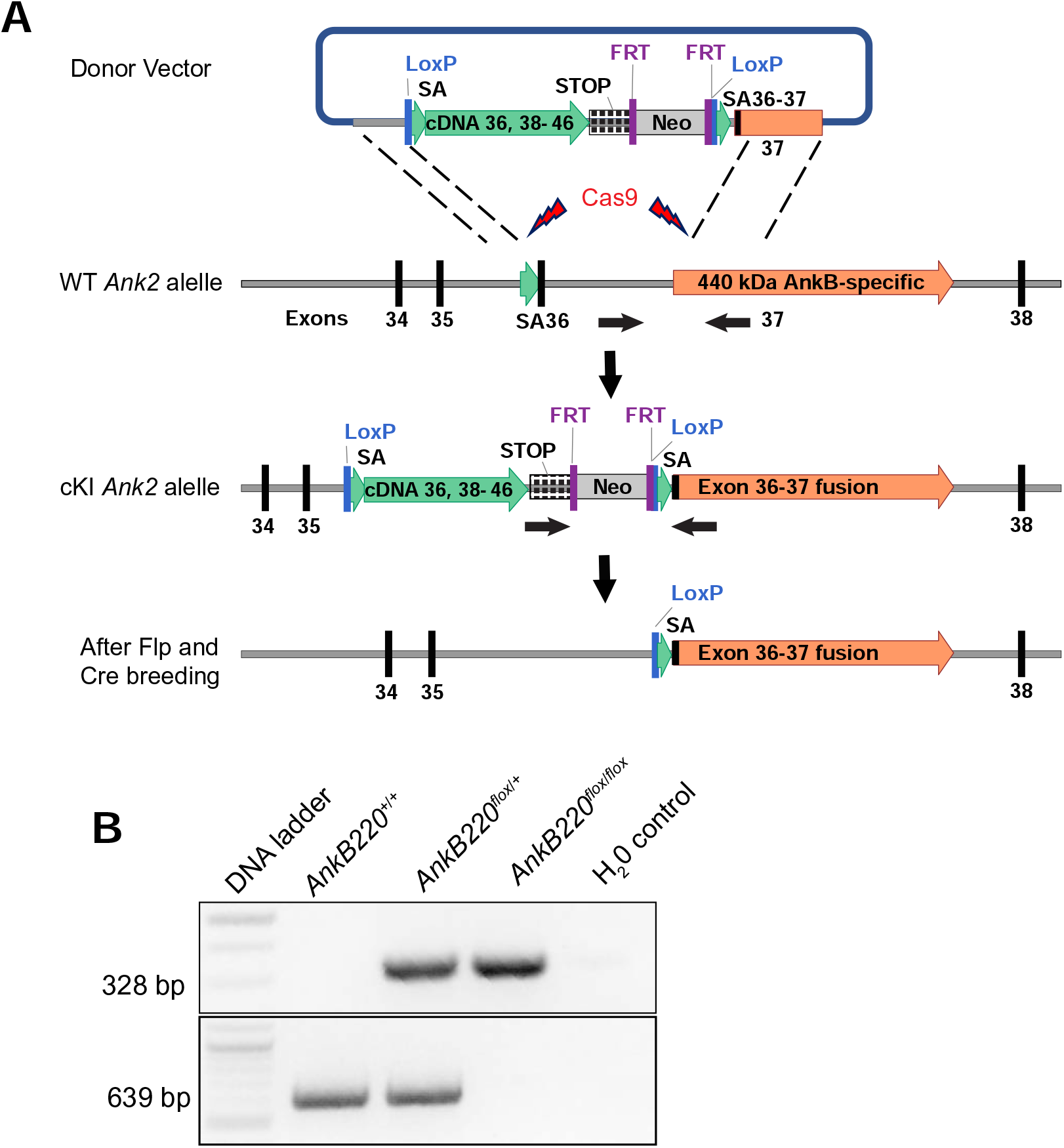
Development of conditional AnkB220 knockout mouse. (**A**) Strategy used to generate a targeting *Ank2* allele (cKI *Ank2*). A donor vector containing a rescue cDNA cassette in which exons 36-37 of *Ank2* were fused to prevent splicing of exon 37 unique to the AnkB440 transcript was introduced into mouse ES cells using CRISPR/Cas9. Transgenic mice bearing the cKI *Ank2* allele were crossed to a mouse line expressing the flippase (FLP) recombinase to eliminate expression of the Neomycin (Neo) selection marker. Breeding to Nestin-Cre resulted in loss of AnkB220 while preserved AnkB440 levels. (**B**) Example of results of genotyping PCR from mouse genomic DNA to identify mice heterozygous or homozygous for the conditional AnkB220^flox^ allele.

**Figure 1—figure supplement 2.**
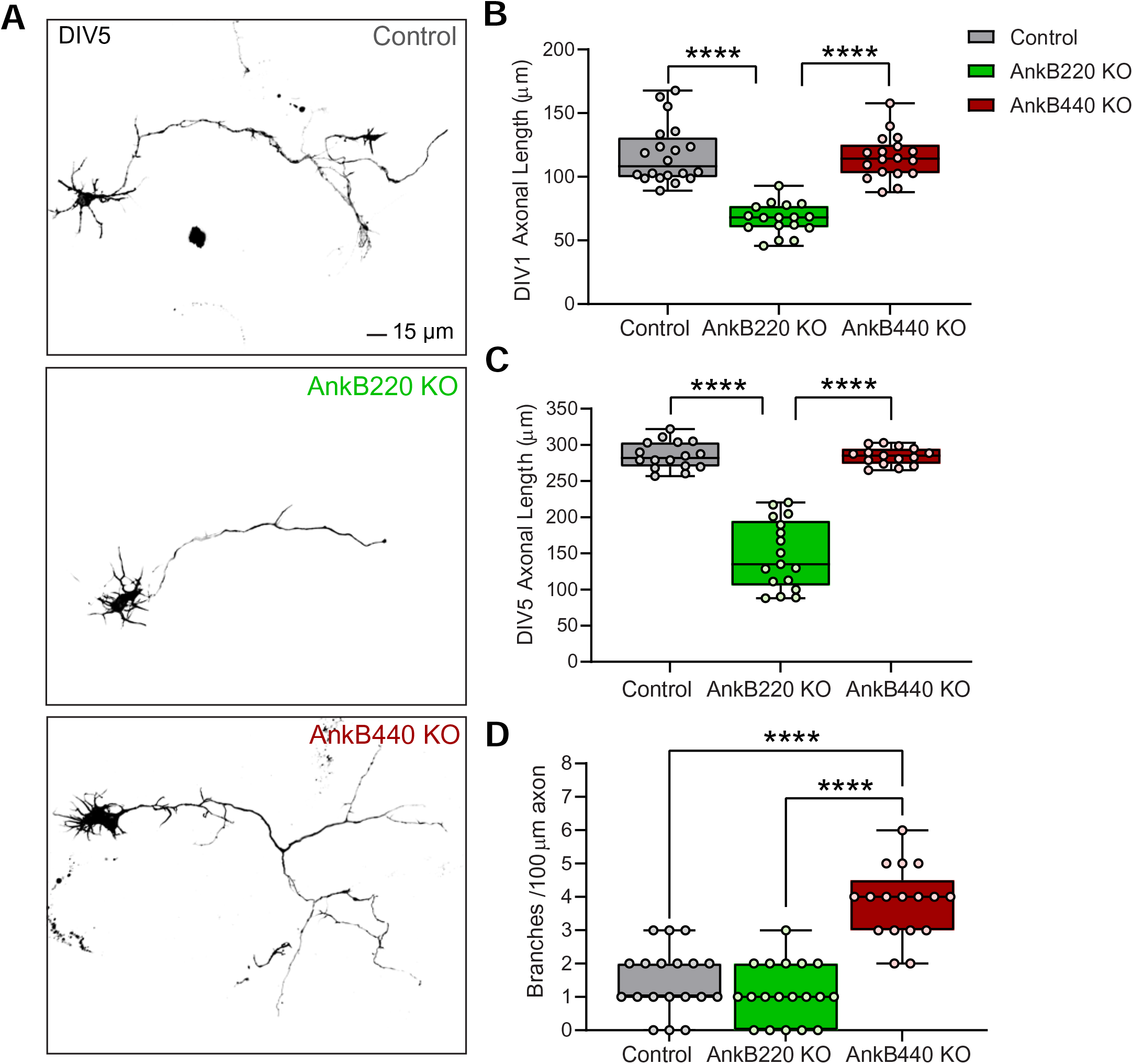
AnkB440 and AnkB220 isoforms exert differential roles in axonal growth and branching. **(A)** Representative images of mScarlet-labeled DIV3 control, AnkB220 KO, and AnkB440 KO cortical neurons. Scale bar, 15 μm. **(B)** Quantification of axonal length at DIV1 of n=20 control, n=17 AnkB220 KO, and n=18 AnkB440 KO mScarlet-labeled neurons from three independent experiments. **(C)** Quantification of axonal length at DIV5 in n=16 control, n=17 AnkB220 KO, and n=15 AnkB440 KO mScarlet-labeled neurons from three independent experiments. **(D)** Quantification of axonal branches per 100 μm of axon of DIV5 (n=19 control, n=19 AnkB220 KO, and n=17 AnkB440 KO) mScarlet-labeled neurons from three independent experiments. The box and whisker plots in **B-D** represent all data points collected arranged from minimum to maximum. One-way ANOVA with Dunnett’s post hoc analysis test for multiple comparisons. ****p < 0.0001.

**Figure 1—figure supplement 3.**
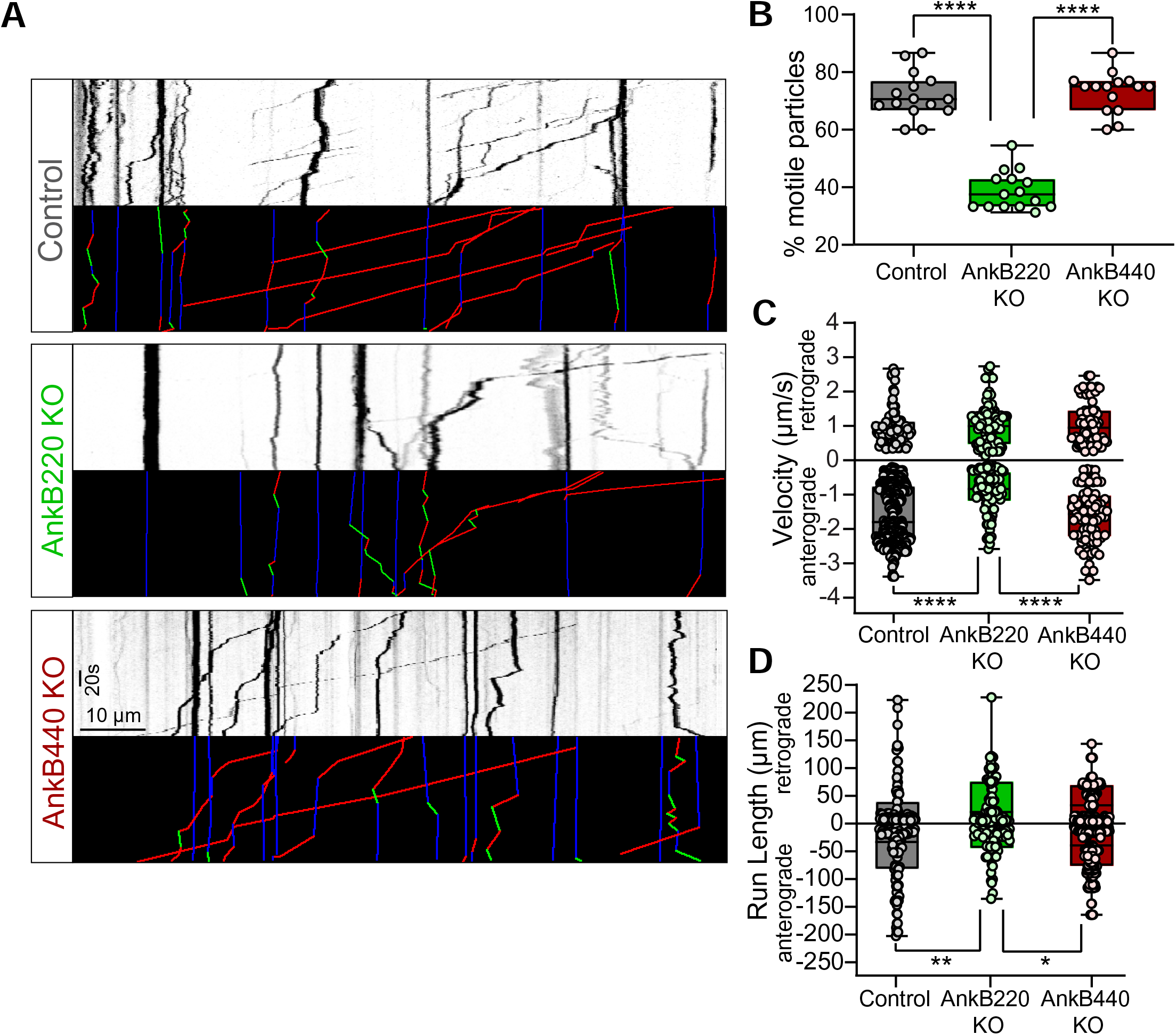
AnkB220, but not AnkB440, promotes axonal organelle transport. (**A**) Kymographs showing motility of GFP-tagged LAMP1 late endosomes/lysosomes at the distal axon of DIV7 neurons from the indicated genotypes. Trajectories in the color-coded kymographs are shown in green for anterograde-moving, red for retrograde-moving, and blue for static cargo (Scale bar, 10 μm and 20 s). (**B**) Percentage of motile GFP-LAMP1 cargo per axon computed from n=15 axons per genotype. (**C**) Anterograde and retrograde velocity of GFP-LAMP1 cargo in axons of indicated genotypes (control: anterograde n=85, retrograde n= 364; AnkB220 KO: anterograde n=87, retrograde n=101; AnkB440 KO: anterograde n=73, retrograde n=76). (**D**) Run length of GFP-LAMP1 cargo in axons of indicated genotypes (control: anterograde n=103, retrograde n= 115; AnkB220 KO: anterograde n=79, retrograde n=65; AnkB440 KO: anterograde n=144, retrograde n=171). The box and whisker plots in **B-D** represent all data points collected arranged from minimum to maximum. One-way ANOVA with Dunnett’s post hoc analysis test for multiple comparisons. ****p < 0.0001, **p < 0.01, *p < 0.05.

**Figure 4—figure supplement 4.**
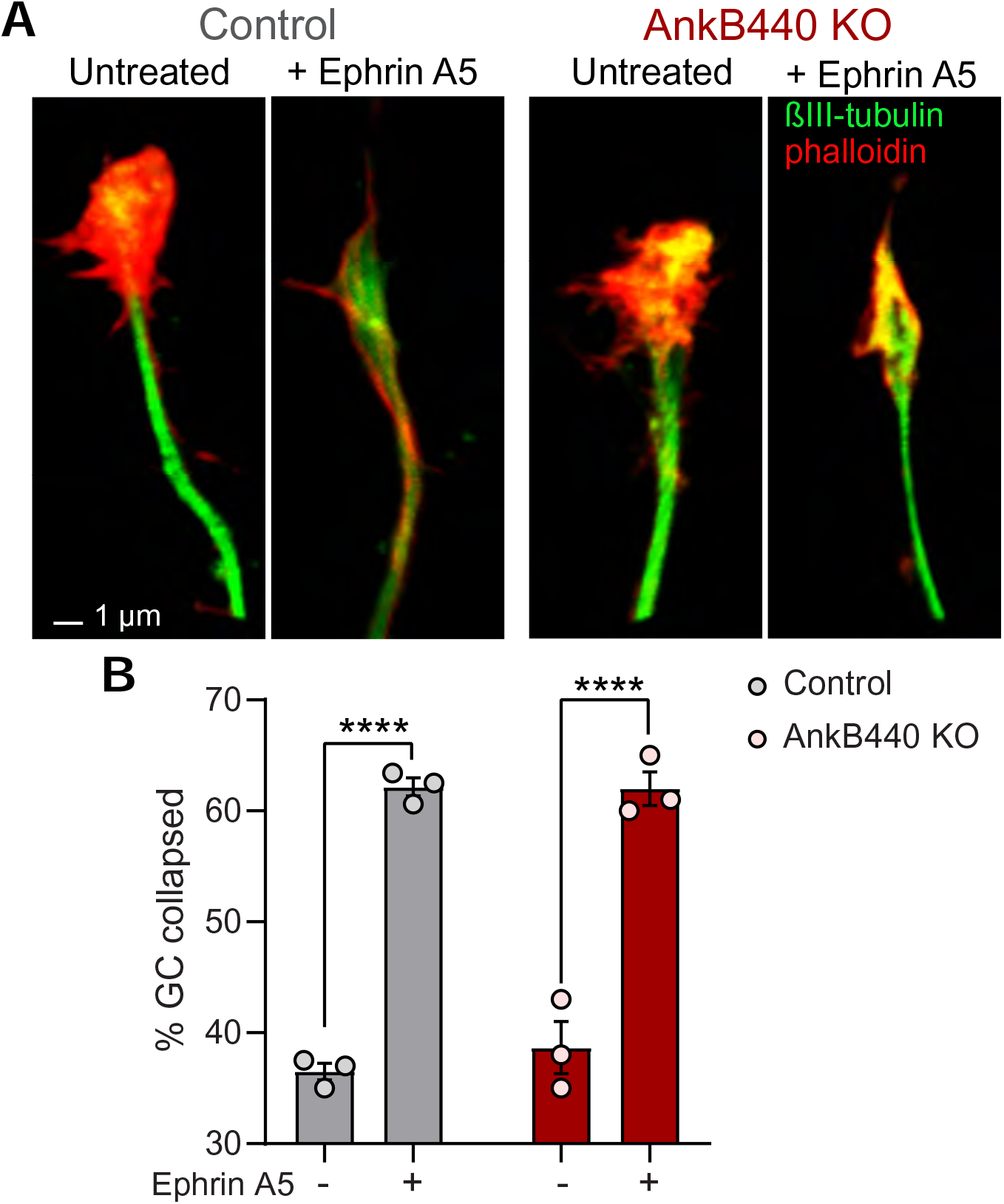
AnkB440 does not enable Ephrin A5-induced GC collapse. (**A**) Images of the distal portion of axons of DIV3 cortical neurons untreated and treated with Ephrin A5 and stained with phalloidin and βIII-tubulin. Scale bar, 1 μm. (**B**) Percentage of GC collapsed before and after Ephrin A5 treatment of control and AnkB440 cortical neurons. Data represent mean ± SEM collected from an average of n=80 GCs/treatment condition/genotype from three independent experiments. Unpaired *t* test. ****p < 0.0001.

**Figure 5—figure supplement 5.**
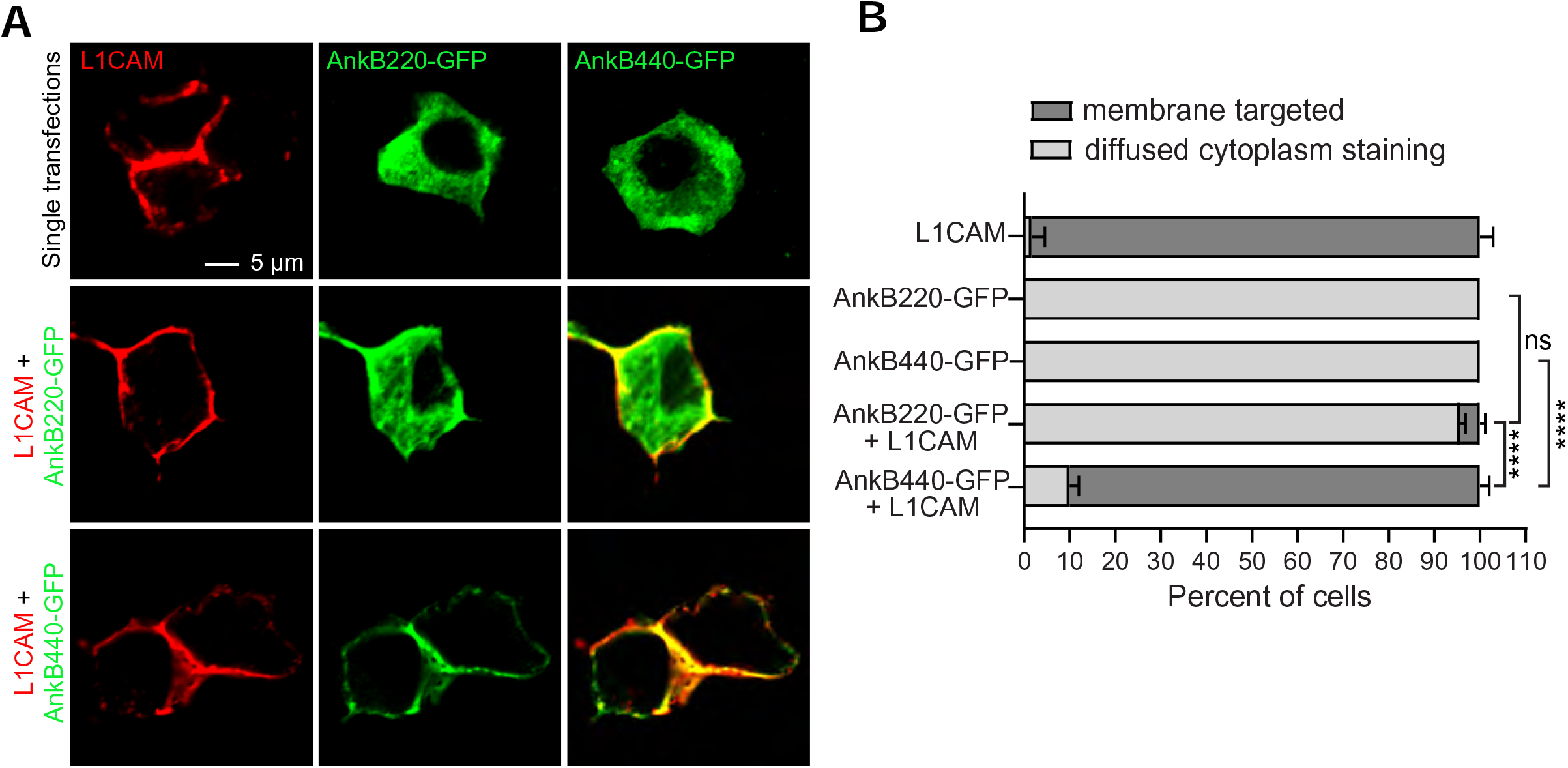
L1CAM recruits AnkB440 to the surface membrane. (**A**) Images of HEK293T cells independently transfected with plasmids encoding L1CAM, AnkB220-GFP, and AnB440-GFP, and L1CAM co-transfected with each of the two GFP-tagged AnkB isoforms. Scale bar, 5 μm. (**B**) Percent of cells in which L1CAM, AnkB220-GFP, or AnkB440-GFP are targeted to the plasma membrane or found diffused throughout the cytoplasm for each experiment. Data represent mean ± SEM collected from an average of n=15 cells/transfection from three independent experiments. One-way ANOVA with Dunnett’s post hoc analysis test for multiple comparisons. ****p < 0.0001, ^ns^p > 0.05.

**Figure 6—figure supplement 6.**
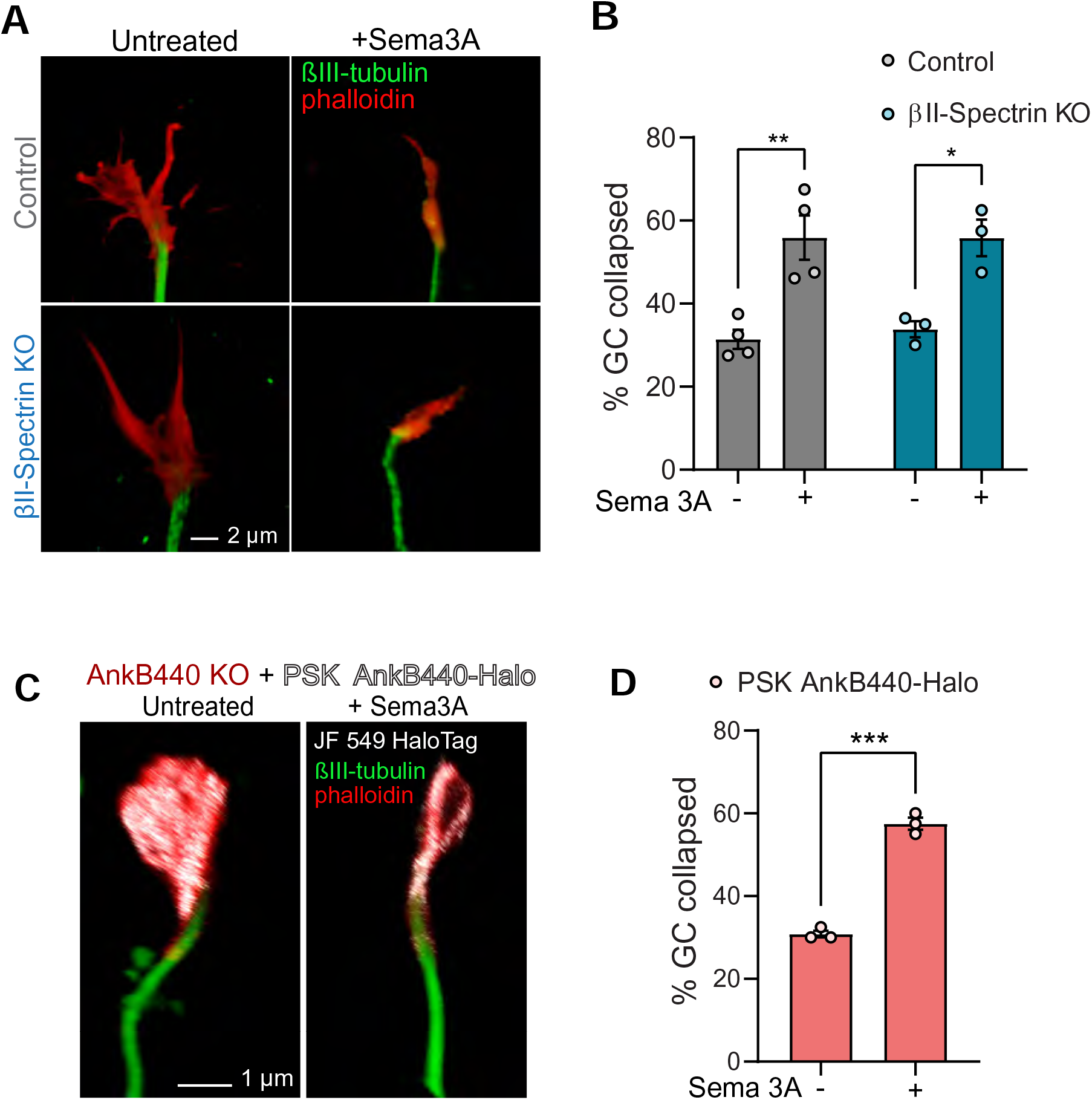
AnkB440 does not require βII-spectrin or binding to microtubules to transduce Sema 3A signaling. (**A**) Images of the distal portion of the main axon of control and βII-pectrin KO DIV3 cortical neurons untreated and treated with Sema 3A and stained with phalloidin and βIII-tubulin. Scale bar, 2 μm. (**B**) Percent of axon GCs that collapse before and after Sema 3A treatment. (**C**) Images of the distal portion of the main axon of DIV3 AnkB 440 KO cortical neurons expressing mutant PSK AnkB440-Halo untreated and treated with Sema 3A. Scale bar, 1 μm. (**B**) Percent main axon GCs that collapse before and after Sema 3A treatment. Data in **B** and **D** represent mean ± SEM collected from an average of n=70-90 GCs/treatment condition/genotype from three independent experiments. Unpaired *t* test. ***p < 0.001, **p < 0.01, *p < 0.05.

**Source data.** Source data contain the original files of the full raw unedited gels or blots and figures with the uncropped gels or blots with the relevant bands clearly labelled.

**Figure 1—Video 1**. **AnkB220 promotes axonal organelle transport.** Timelapse images illustrate the motility of GFP-tagged LAMP1 late endosomes/lysosomes at the distal axon of DIV7 cortical neurons from the indicated genotypes. Movies are played at 15 frames/second. Scale bar, 10 μm.

**Figure 5—Video 2**. **The AnkB440-L1CAM complex promotes GC collapse induced by Sema 3A.** Timelapse images illustrate changes in F-actin dynamics at the GC of collateral axon branches and fillopodia of DIV3 cortical neurons in response to Sema 3A. Movies are played at 15 frames/second. Scale bar, 5 μm.

